# Targeting soluble immunoglobulins ameliorates idiopathic multicentric Castleman disease in a mouse model

**DOI:** 10.64898/2026.07.26.740764

**Authors:** David C. Uhlfelder, David Andruszewski, Leon R. Amelong, Helena S. Schäfer, Mia Hüttenschmidt, Michaela Blanfeld, Carsten Schelmbauer, Zeynep Ergün, Aysan Poursadegh Zonouzi, Anne Hausen, Stefanie Zimmer, Matthias M. Gaida, Saskia von Ungern-Sternberg, Kerstin Jurk, F. Thomas Wunderlich, Gabriel Azevedo Publio, Daniele C. Nascimento, Jose Carlos Alves-Filho, Thomas Korn, Ari Waisman, Ilgiz A. Mufazalov

## Abstract

Idiopathic multicentric Castleman disease (iMCD) is a rare lymphoproliferative disorder that affects lymph nodes at multiple sites and can progress to life-threatening organ dysfunction. While curative therapies remain unavailable, largely due to a critical gap in understanding the disease’s underlying etiology, the cytokine interleukin-6 (IL-6) is recognized as a major pathogenic driver, particularly in iMCD patients with plasmacytic lymphadenopathy. Here, we demonstrate the presence of Foxp3-expressing cells alongside plasma cells within germinal centers of lymph nodes from such iMCD patients. Based on this observation, we directed IL-6 overexpression specifically to Foxp3^+^ Treg cells and established a novel preclinical mouse model of iMCD. This targeted approach successfully recapitulated the key characteristics of iMCD in mice, including multifocal lymphadenopathy, splenomegaly, anemia, thrombocytopenia, marked plasmacytosis, and mortality in young adults. Advanced disease was characterized by renal pathology, including proteinuria and elevated creatinine levels. In this model, IL-6 directly promoted plasmacytosis, resulting in profound IgG1 hyperimmunoglobulinemia. Genetic ablation of soluble immunoglobulin production ameliorated iMCD symptoms, prevented renal dysfunction, and significantly extended survival. Our findings identify IL-6-driven IgG1 plasmacytosis as a critical pathogenic factor in iMCD, shedding light on the mechanisms underlying its development.

**Key points:**

1. Transgenic IL-6 expression in Foxp3^+^ cells models iMCD in mice and drives lethal IgG1 plasmacytosis
2. Loss of soluble immunoglobulins prevented renal impairment and prolonged survival in iMCD-like mice

## Introduction

Multicentric Castleman disease (MCD) is a rare lymphoproliferative disorder characterized by generalized lymphadenopathy. Idiopathic MCD (iMCD) is the HHV-8-negative subtype with unknown etiology^1,2^. Severe iMCD is associated with life-threatening multiorgan dysfunction^3^. Interleukin-6 (IL-6) is considered a major pathogenic driver of iMCD, and the only approved therapies target IL-6 signaling, either through the IL-6 receptor (tocilizumab, approved in Japan^4^) or IL-6 itself (siltuximab, approved in the USA and the European Union^5^). Although IL-6 blockade provides substantial clinical benefit in some patients, many respond inadequately or eventually relapse. Additional pathways have been implicated in iMCD through biomarker studies and beneficial responses to targeted therapies, including CXCL13^6–9^ and VEGF^10^, TNF^11^, IL-1^12–17^, mTOR^18–20^, the proteasome^21^, Janus kinsase^22^, and Bruton’s tyrosine kinase^23^. Nevertheless, the disease often carries a poor prognosis^24^.

To accelerate the development of a cure, the Castleman Disease Collaborative Network (CDCN) prioritized several research pipelines aiming at understanding the mechanisms of iMCD development^25^. One of the key obstacles hindering progress in iMCD research is the limited availability of patient biospecimens due to the rarity of the disease. Preclinical mouse models therefore represent a valuable alternative for mechanistic studies. Here, we report a novel murine iMCD model that fulfills the most recent consensus diagnostic criteria for iMCD established by the CDCN in 2017^26^. To generate this model, we used the Cre-loxP system to induce IL-6 overexpression in a defined cell type^27^. We selected Foxp3-Cre based on histological analyses of human lymph nodes, which revealed FOXP3⁺ cells adjacent to plasma cells in iMCD patients, particularly those with plasmacytic lymphadenopathy. Using this strategy, we generated mice with Foxp3-Cre-mediated IL-6 overexpression (IL-6^Fp3-OE^ mice), which faithfully recapitulated key features of iMCD.

IL-6^Fp3-OE^ mice developed multifocal lymphadenopathy, splenomegaly, anemia, thrombocytopenia, plasmacytosis, and hyperimmunoglobulinemia with complete penetrance. Plasma cells retained surface IL-6 receptor expression and exhibited downstream STAT3 phosphorylation. End-stage disease was critically dependent on excessive immunoglobulin production, particularly IgG1, resulting in proteinuria and elevated serum creatinine indicative of renal dysfunction. Genetic ablation of soluble immunoglobulin secretion, including IgG1, significantly prolonged maximum survival from 14 to 30 weeks in IL-6-overexpressing mice. Collectively, our findings provide experimental evidence that IL-6-driven IgG1 plasmacytosis is a key pathogenic mechanism underlying life-threatening iMCD-like disease.

## Methods

### Human samples

Lymph node samples from patients with HHV-8-negative Castleman disease (n=7) were collected between 2002 and 2021 and provided by the Tissue Bank of the University Medical Center Mainz in accordance with institutional biobank regulations and approval by the Ethics Committee of the University Medical Center Mainz. The retrospective histological analysis was approved by the Ethics Committee of the Landesärztekammer Rheinland-Pfalz.

Paraffin-embedded samples were sectioned, stained with hematoxylin (Dako EnVision FLEX Mayer’s Hematoxylin, S3309), and incubated overnight with an anti-FOXP3 antibody (Abcam, ab20034). Detection was performed using Dako EnVision FLEX DAB+ Chromogen (DM827).

### Mice

Rosa26^IL-6(stop/stop)^ mice carrying a Cre-loxP-dependent murine *Il6* cDNA transgene inserted into the Rosa26 locus were previously generated by us^27,28^. These mice were crossed with Foxp3-Cre mice^29^ to generate experimental IL-6^Fp3-OE^ mice with IL-6 overexpression in Foxp3⁺ regulatory T cells (**Supplemental Figure 1**). IL-6^Fp3-OE^ mice were further crossed with IgH^μγ1/μγ1^ mice^30^, which carry a mutated IgH locus that prevents secretion of soluble immunoglobulins. *Il6ra* flox mice^31^ were crossed with CD11c-Cre mice^32^ to generate IL-6Rα knockout mice, as described previously^27^. Detailed breeding strategies are provided in the Supplemental methods. All animal experiments were approved by the Landesuntersuchungsamt Rheinland-Pfalz and performed in accordance with the German Animal Welfare Act.

### Murine blood preparation

For cytokine and creatinine measurements, blood was collected into serum gel tubes (Sarstedt™, 1.1 mL), allowed to clot for 30 minutes at room temperature, centrifuged (10,000 × g, 10 minutes), and serum was stored at −80°C. ELISA procedures and creatinine measurements are described in the Supplemental methods. For complete blood counts, blood was collected into K3 EDTA tubes (Sarstedt™, 1.3 mL) and analyzed using a Sysmex KX-21N hematology analyzer. For flow cytometry, erythrocytes were lysed with ACK lysis buffer.

### Urine protein measurement

Proteinuria was assessed using 20 μL of midstream urine applied to a urine dipstick (Roche Combur® HC 5 Test) for 60 seconds according to the manufacturer’s instructions.

### Mini-endoscopy

Colonic inflammation was assessed by high-resolution mini-endoscopy (Karl Storz SE & Co. AG) in anesthetized mice and scored for stool consistency, translucency, granularity, vascularity, and fibrin, as previously described^33^.

### Preparation of murine single-cell suspensions

Single-cell suspensions from lymph nodes, spleens, and bone marrow were obtained by mechanical dissociation through 40 μm strainers into phosphate-buffered saline (PBS) supplemented with 2% fetal calf serum (FCS) and kept on ice. For B-cell transfer experiments, CD19⁺ B cells were enriched using magnetic beads (Miltenyi Biotec), as described in the Supplemental methods. Colonic lamina propria lymphocytes (LPLs) were isolated by mechanical tissue dissociation and enzymatic digestion without Percoll gradient centrifugation, as previously described^34^. Kidney immune cells were isolated after cardiac perfusion and mechanical dissociation, also without a Percoll gradient, as previously described^35^.

Cell counts were determined using a LUNA-II Automated Cell Counter (Logos Biosystems, South Korea). Flow cytometry procedures are described in the Supplemental methods.

### Anti-nuclear antibody (ANA) detection in murine serum

ANA were detected on HEp-2 biochips (EuroImmun, Cat. No. FB 1510-1010-1) according to the manufacturer’s instructions. Bound antibodies were visualized using AlexaFluor 594-conjugated anti-mouse IgG1 (Invitrogen) with DAPI counterstaining and imaged at 20× magnification using an Olympus IX-81 microscope.

### Murine Histology

Inguinal lymph nodes, spleen, kidney, ear skin, and colon were fixed in ROTI® Histofix (Carl Roth GmbH + Co. KG) for at least 24 hours, paraffin embedded, and sectioned at 2 μm using a Hyrax M55 microtome (Zeiss). Hematoxylin and eosin (H&E; Dako, CS70030-2 and CS701), periodic acid–Schiff (PAS; Carl Roth, 3257.1; Supelco, 1.06268.0250; Sigma-Aldrich, 1.05175.2500), and Ki67 immunohistochemical staining (Bethyl, ICH-00375) were performed according to the manufacturers’ instructions. Whole-slide images were acquired using an Aperio AT2 slide scanner (Leica Biosystems).

For renal immunofluorescence, kidneys were snap-frozen, embedded in Tissue-Tek® O.C.T. compound (Sakura Finetek), sectioned at 4 μm using a MEV SLEE cryostat, and fixed in ice-cold acetone for 10 minutes. Sections were blocked for 1 hour in TBST containing 10% goat serum and 2% bovine serum albumin (BSA), followed by overnight incubation at 4°C with rat anti-mouse IgG1 antibody (clone A85-1, BD Pharmingen). After washing, sections were incubated with CF555-conjugated goat anti-rat IgG secondary antibody (Sigma-Aldrich), mounted in Vectashield® Antifade Mounting Medium containing DAPI (Vector Laboratories), and imaged using a Keyence BZ-X810 fluorescence microscope.

### Joint measurements

The diameters of ankle and wrist joints were measured using a digital caliper with a resolution of 0.01 mm (Vogel, Germany).

### Statistical analysis

Statistical analysis and graphical representation were performed with Prism 10 software (GraphPad). Statistical significance was calculated using unpaired two-tailed t test, one-way ANOVA, and Gehan–Breslow–Wilcoxon test for Kaplan–Meier survival curves. P values < 0.05, < 0.01, and < 0.001 were indicated by *, **, and ***, respectively.

## Results

### FOXP3-positive cells are abundantly present alongside plasma cells in the lymph nodes of iMCD-IPL patients

Originally, IL-6 was described as a B-cell stimulatory factor, and its role in plasma cell induction has been demonstrated in multiple studies^36^. Likewise, IL-6 is a key driver of iMCD pathology, particularly in the idiopathic plasmacytic lymphadenopathy (iMCD-IPL) subtype, which is characterized by marked plasmacytosis^37^. However, the cellular source of pathogenic IL-6 in iMCD remains unclear, likely due to the poorly defined local microenvironment where plasma cells reside. Previous studies have suggested that Foxp3⁺ cells are located in close proximity to plasma cells in mice^38^.

To determine whether FOXP3^+^ cells can also be detected alongside plasma cells in clinical settings, we analyzed lymph node samples from iMCD patients using immunohistochemistry. We found abundant FOXP3 expression in the germinal centers of lymph nodes from patients with plasmacytic iMCD subtype. In contrast, iMCD patients with hyaline-vascular histopathology, as well as those with unicentric Castleman disease (UCD), exhibited very few plasma cells and FOXP3-positive cells in their lymph nodes (**Figure 1A-B, Supplemental Figure 1**). We concluded that the abundance of FOXP3^+^ cells in germinal centers of iMCD-IPL lymph nodes could be leveraged to establish a faithful mouse model of iMCD.

**Figure 1.**
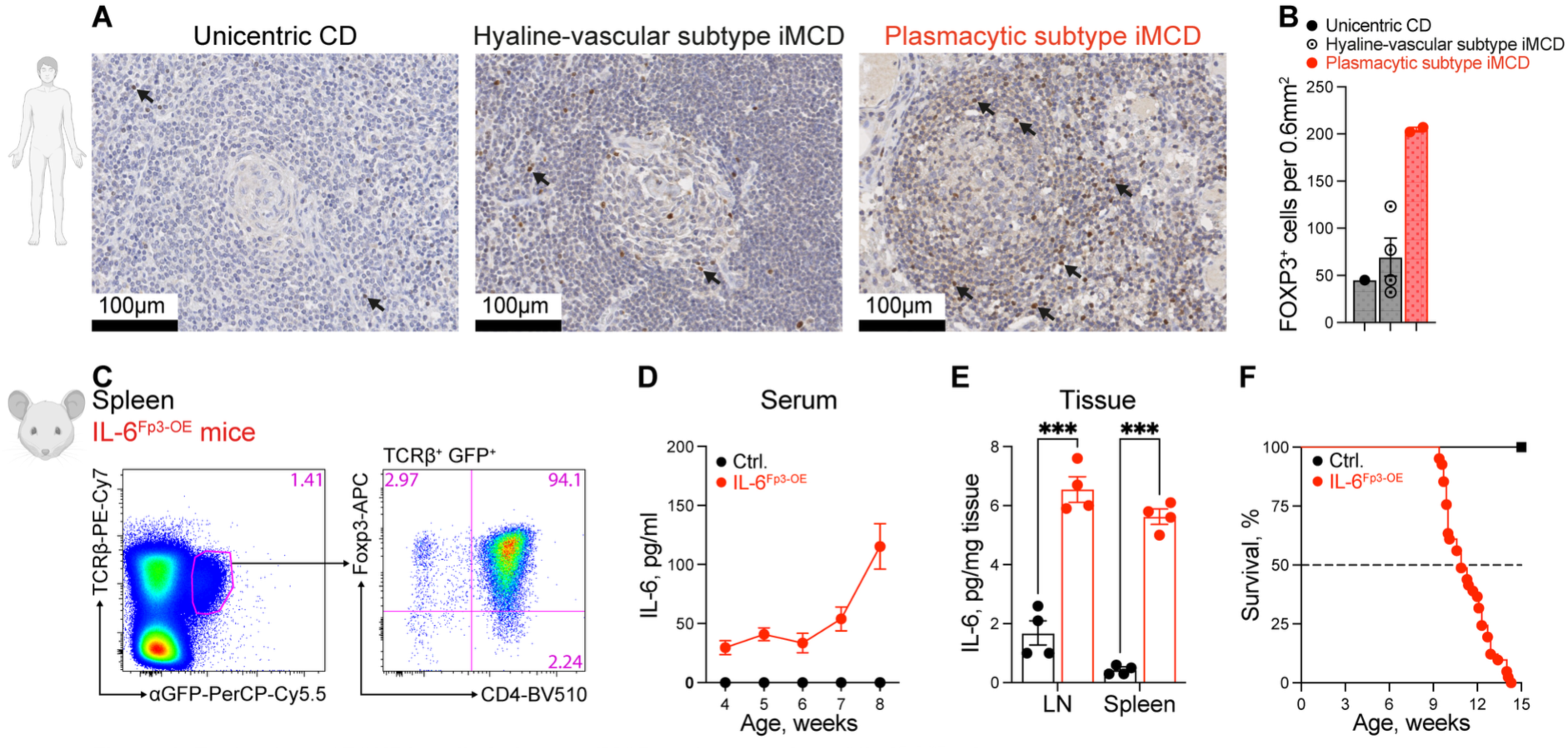
Evaluation of FOXP3^+^ cells in human iMCD and generation of IL-6^Fp3-OE^ mice. **A** – Analysis of FOXP3 expression (brown) in germinal centers of lymph nodes from patients with unicentric Castleman disease (UCD), hyaline-vascular iMCD subtype, and plasmacytic iMCD subtype. Scale bars indicate 100 μm. Black arrows indicate FOXP3-positive cells. **B** – Quantification of FOXP3^+^ cells in germinal centers per 0.6 mm² in the lymph nodes of UCD and iMCD patients. **C** – FACS analysis of eGFP expression in splenocytes from IL-6^Fp3-OE^ mice. Cells were stained with anti-GFP antibodies and analyzed for co-expression of TCRβ, CD4 and Foxp3. **D** – IL-6 levels in sera at the indicated ages, measured by ELISA. n=5 control mice; n=5 IL-6^Fp3-OE^ mice. **E** – IL-6 levels in lymph nodes and spleens of 7–8-week-old mice, measured by ELISA. **F** – Kaplan-Meier survival curves of control (n=10) and IL-6^Fp3-OE^ (n=41) mice. Graphs (B, D, E) show mean ± SEM. Each dot represents an individual mouse. P values in (E) were calculated using unpaired Student’s t-test and are indicated as follows: *** (p < 0.001).

### Generation of the IL-6^Fp3-OE^ mouse strain to model iMCD

Based on our observation that FOXP3-positive cells are abundant in germinal centers adjacent to plasma cells in patients with the plasmacytic iMCD subtype, we hypothesized that IL-6 expression by Foxp3-expressing cells could recapitulate iMCD in mice. To test this, we used transgenic mice carrying a Cre-loxP-dependent murine *Il6* cDNA transgene, enabling IL-6 overexpression (OE) following Cre-mediated excision of a stop cassette^27,28^. We crossed these mice with Foxp3-Cre (Fp3) mice to generate IL-6^Fp3-OE^ mice, in which IL-6 is overexpressed specifically in Foxp3^+^ regulatory T (Treg) cells, along with an eGFP reporter (**Supplemental Figure 2**). As expected, the vast majority of eGFP-positive cells in IL-6^Fp3-OE^ mice were CD4⁺Foxp3⁺ Treg cells, reflecting Foxp3-Cre activity (**Figure 1C**). Activation of the *Il6* transgene resulted in elevated serum IL-6 levels that progressively increased with age (**Figure 1D**). Similarly, IL-6 levels were elevated in the lymph nodes and spleens of IL-6^Fp3^ ^OE^ mice (**Figure 1E**).

High levels of IL-6 are lethal in both humans and mice^39^. Consistent with our previous mouse models of IL-6 overexpression^27,28^, IL-6^Fp3-OE^ mice also exhibited complete mortality. IL-6^Fp3-OE^ mice died spontaneously between 9 and 14 weeks of age or were euthanized upon reaching humane endpoints (**Figure 1F**).

### IL-6^Fp3-OE^ mice develop generalized lymphadenopathy with defined plasmacytic histopathology – two major criteria in iMCD diagnosis

After generating the IL-6^Fp3-OE^ mouse model, we assessed whether it fulfilled the diagnostic criteria for iMCD. IL-6^Fp3-OE^ mice developed enlarged lymph nodes at multiple anatomical sites starting at 4 weeks of age, with progressive enlargement over time (**Figure 2A**). This generalized lymphadenopathy represents one of the two major diagnostic criteria for iMCD^26^. Histological analysis further revealed pronounced hypervascularization (**Figure 2B**) and plasmacytosis (**Figure 2C**), fulfilling the second major diagnostic criterion.

**Figure 2.**
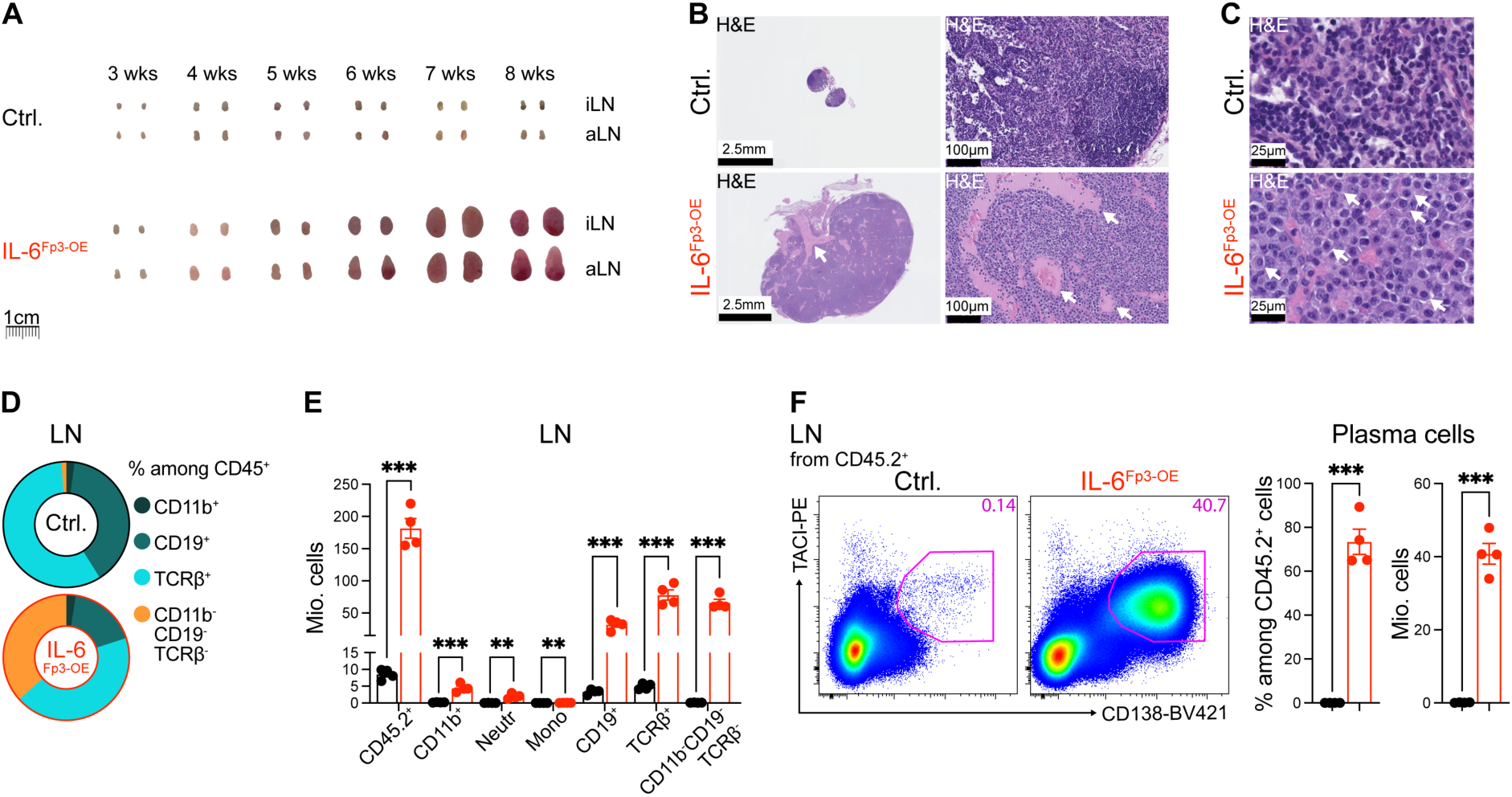
Analysis of major diagnostic criteria for iMCD in IL-6^Fp3-OE^ mice. **A** – Macroscopic examination of axillary lymph nodes (aLN) and inguinal lymph nodes (iLN) at the indicated ages. **B** – Histological analysis (H&E staining) of inguinal lymph nodes. Scale bars: 2.5mm (left column) and 100μm (right column). White arrows indicate hypervascularization. n=4 mice per genotype, 9.5 weeks old. **C** – Histological analysis (H&E staining) of inguinal lymph nodes. Scale bars: 25μm. White arrows highlight representative plasma cells. n=4 mice per genotype, 8 weeks old. **D** – Frequency distribution of indicated immune cell subsets in pooled inguinal, axial, and brachial lymph nodes based on flow cytometry analysis. n=4 mice per genotype, 8 weeks old. **E** – Absolute numbers of immune cells in lymph nodes shown in (D). **F** – FACS analysis of CD138^+^TACI^+^ plasma cells in lymph nodes. n=4 mice per genotype, 8 weeks old. Graphs (E, F) show mean ± SEM. Each dot represents an individual mouse. P values in (E, F) were calculated using unpaired Student’s t-test and are indicated as follows: ** (p < 0.01), *** (p < 0.001).

To characterize the immune landscape of affected lymph nodes, we analyzed their cellular composition by flow cytometry. IL-6^Fp3-OE^ mice exhibited a marked increase in CD45⁺ leukocytes, including CD11b⁺ myeloid cells, CD19⁺ B cells, and TCRβ⁺ T cells. The most prominent expansion occurred within the CD11b^-^CD19^-^TCRβ^-^ population (**Figure 2D-E**). Further analysis identified these cells predominantly as CD138⁺TACI⁺ plasma cells, which were dramatically expanded in the lymph nodes of IL-6^Fp3-OE^ mice (**Figure 2F**).

### IL-6^Fp3-OE^ mice exhibit several minor diagnostic criteria of iMCD, accompanied by renal dysfunction

After detecting both major iMCD criteria in IL-6^Fp3-OE^ mice, we proceeded to analyze the minor criteria, of which at least two out of eleven are sufficient to establish iMCD diagnosis^26^. First, splenomegaly, one of the minor diagnostic criteria for iMCD^26^, was a consistent finding in IL-6^Fp3^ ^OE^ mice (**Figure 3A**). Flow cytometric analysis revealed an increase in the numbers of CD45⁺ splenocytes, including CD11b⁺, CD19⁺, and TCRβ⁺ cells, as well as immune cells negative for these markers (**Figure 3B**). Similar to the lymph nodes, the spleens of IL-6^Fp3^ ^OE^ mice accumulated large numbers of plasma cells (**Figure 3C**), which was confirmed by histological analysis (**Figure 3D**).

**Figure 3.**
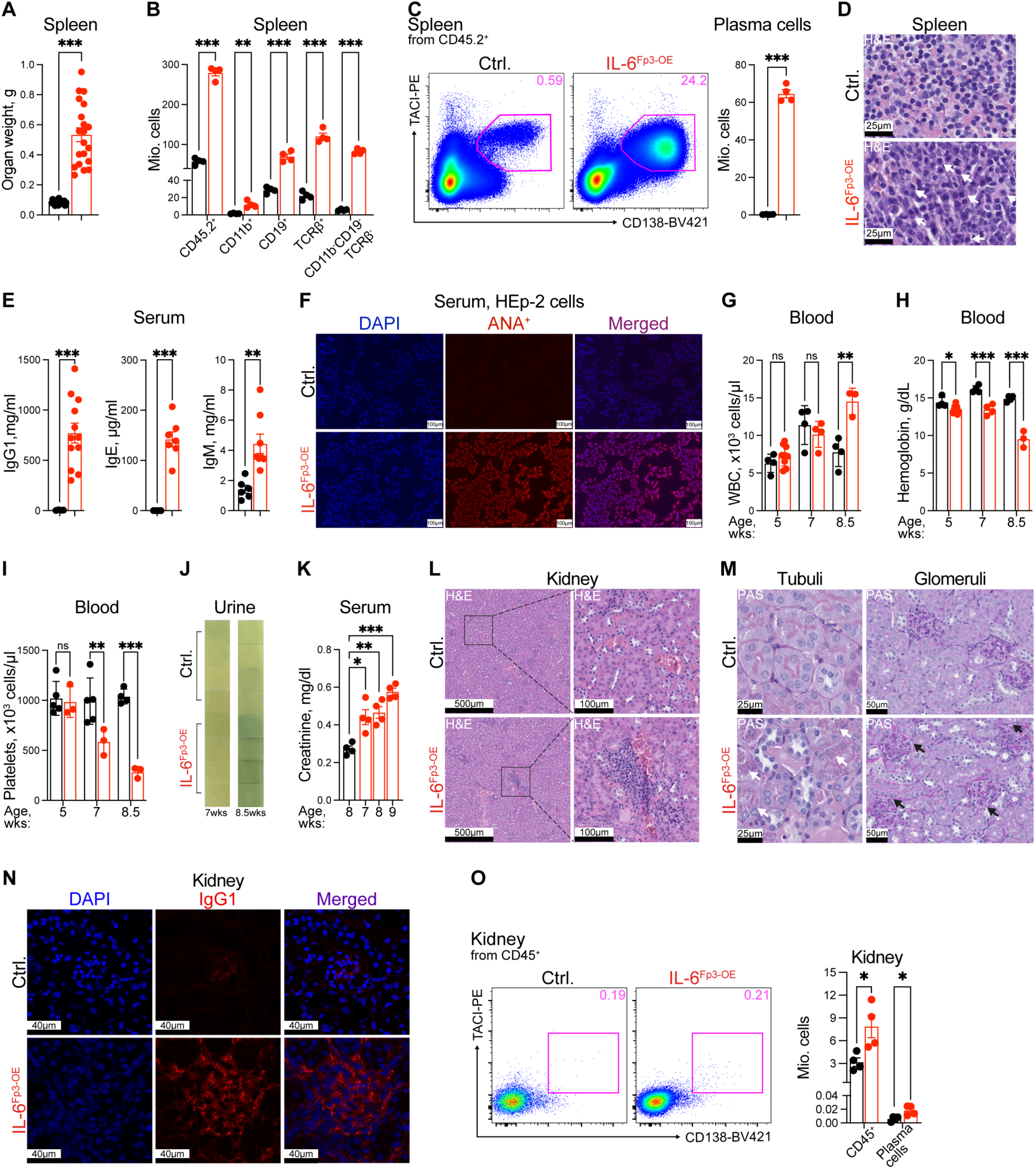
Analysis of minor diagnostic criteria for iMCD and kidney dysfunction in IL-6^Fp3-OE^ mice. **A** – Spleen weight analysis of 8-weeks-old mice. **B** – Absolute numbers of immune cells in the spleen based on FACS analysis. n=4 mice per genotype, 8 weeks old. **C** – FACS analysis of CD138^+^TACI^+^ plasma cells in the spleen. n=4 mice per genotype, 8 weeks old. **D** – Histological analysis (H&E staining) of spleen. Scale bars: 25 μm. White arrows highlight representative plasma cells. n=4 mice per genotype, 8 weeks old. **E** – IgG1, IgE, and IgM levels in the serum of 8- to 9-weeks-old mice measured by ELISA. **F** – Analysis of antinuclear antibodies (ANA) in the serum of 8-week-old control and IL-6^Fp3-OE^ mice, detected by immunofluorescence on HEp-2 cells. IgG1 autoantibodies (red) and nuclei (DAPI, blue). Scale bar indicates 100 μm. **G** – Analysis of white blood cells (WBC) at indicated age. **H** – Hemoglobin measurements at indicated age. **I** – Platelet counts at indicated age. **J** – Urine proteins measured by dipsticks (Roche Combur Test® HC) at indicated age. **K** – Creatinine levels in serum measured by a colorimetric assay at indicated age. **L** – Histological analysis (H&E staining) of kidneys depict vasculature with mononuclear immune cell infiltration. Scale bars: 500 μm (left panels) and 100 μm (right panels). n=6 mice per genotype, 9.5 weeks old. **M** – Histological analysis (Periodic acid-Schiff (PAS)) staining of kidney sections. White arrows indicate granular inclusions in the tubular epithelial cells, black arrows indicate thickened Bowman capsule. Scale bars: 25 μm (left panels) and 50 μm (right panels). **N** – Immunofluorescence staining of kidneys for IgG1 immunocomplex deposition. IgG1 (red); nuclei stained with DAPI (blue). Scale bars: 25 μm. n=3 mice per genotype, 8 weeks old. **O** – FACS analysis of plasma cells in kidney. Graphs (A, B, C, E, G, H, I, K, O) show mean ± SEM. Each dot represents an individual mouse. P values in (A, B, C, F, G, H, O, E, J) were calculated using unpaired Student’s t-test; and are indicated as follows: * (p < 0.05), ** (p < 0.01), *** (p < 0.001); ns – not significant (p > 0.05).

Hypergammaglobulinemia is a minor diagnostic criterion for iMCD, whereas elevated IgE is considered a supportive finding^26^. Accordingly, IL-6^Fp3-OE^ mice exhibited markedly increased serum levels of IgG1, IgE, and IgM (**Figure 3E**). Among these, IgG1 was the most strongly elevated isotype. Furthermore, sera from IL-6^Fp3-OE^ mice showed strong self-reactive anti-nuclear antibody (ANA) IgG1 staining (**Figure 3F**), consistent with reports that autoantibodies are present in approximately 30% of patients with iMCD^2^.

Altered hematologic parameters constitute additional minor diagnostic criteria for iMCD^26^. Accordingly, adolescent and young adult IL-6^Fp3-OE^ mice developed peripheral leukocytosis, reflected by increased white blood cell counts, together with reduced hemoglobin levels indicative of anemia (**Figure 3G-H**). Platelet counts were also markedly reduced in IL-6^Fp3-OE^ mice compared with age-matched controls from 7 weeks of age onward (**Figure 3I**).

Renal dysfunction is common in iMCD, with up to 48% of patients developing acute kidney injury and 16% progressing to chronic renal insufficiency^40^. Proteinuria and reduced estimated glomerular filtration rate (eGFR) are recognized minor diagnostic criteria^26^. Accordingly, adult IL-6^Fp3-OE^ mice developed proteinuria indicative of glomerular injury (**Figure 3J**). In addition, serum creatinine levels progressively increased with age, consistent with declining renal function (**Figure 3K**).

To investigate the underlying causes of renal dysfunction, we examined kidney sections and detected interstitial neovascularization and perivascular mononuclear cell infiltration in IL-6^Fp3-OE^ mice (**Figure 3L**). Periodic acid-Schiff (PAS) staining further revealed granular PAS-positive inclusions within renal tubules and thickening of Bowman’s capsule, indicative of pathological parenchymal changes of the kidney (**Figure 3M)**.

Given the markedly elevated serum IgG1 levels in IL-6^Fp3-OE^ mice, we hypothesized that renal dysfunction may result from immune complex deposition. Indeed, immunofluorescence staining demonstrated prominent IgG1 deposition in the kidneys of IL-6^Fp3-OE^ mice (**Figure 3N**). In contrast, CD138⁺TACI⁺ plasma cells represented less than 1% of renal immune cells and were only minimally increased despite substantial CD45⁺ leukocyte infiltration (**Figure 3O**). These findings suggest that renal dysfunction in IL-6^Fp3-OE^ mice is driven primarily by IgG1 immune complex deposition rather than local plasma cell accumulation, consistent with observations in human iMCD^41,42^.

Collectively, IL-6^Fp3-OE^ mice fulfilled both major and five minor diagnostic criteria for iMCD, including splenomegaly, hypergammaglobulinemia, anemia, thrombocytopenia, and renal dysfunction with proteinuria. Elevated serum IgE levels and structural alterations of the renal tubules and glomeruli provided additional features consistent with human iMCD.

### IL-6^Fp3-OE^ mice do not meet exclusion criteria for iMCD diagnosis

iMCD overlaps with several autoimmune and malignant disorders that must be excluded before diagnosis^26^. To determine whether IL-6 overexpression disrupted peripheral tolerance and induced barrier-organ autoimmunity, we examined the skin and gut, tissues typically affected in scurfy-like autoimmune disease^43,44^. Unlike scurfy mice, IL-6^Fp3-OE^ mice showed no overt skin pathology (**Figure 4A**). Histological analysis of ear skin revealed no dermal thickening or inflammatory infiltrates, consistent with the absence of Ki67⁺ immune cell accumulation (**Figure 4B**). Likewise, IL-6^Fp3-OE^ mice gained weight similarly to controls (**Figure 4C**), and mini-endoscopy, immune cell analysis of the colonic lamina propria, and histological examination showed no evidence of colonic inflammation (**Figure 4D, Supplemental Figure 3**). Together, these findings indicate that IL-6^Fp3-OE^ mice do not develop systemic scurfy-like autoimmunity^43,44^.

**Figure 4.**
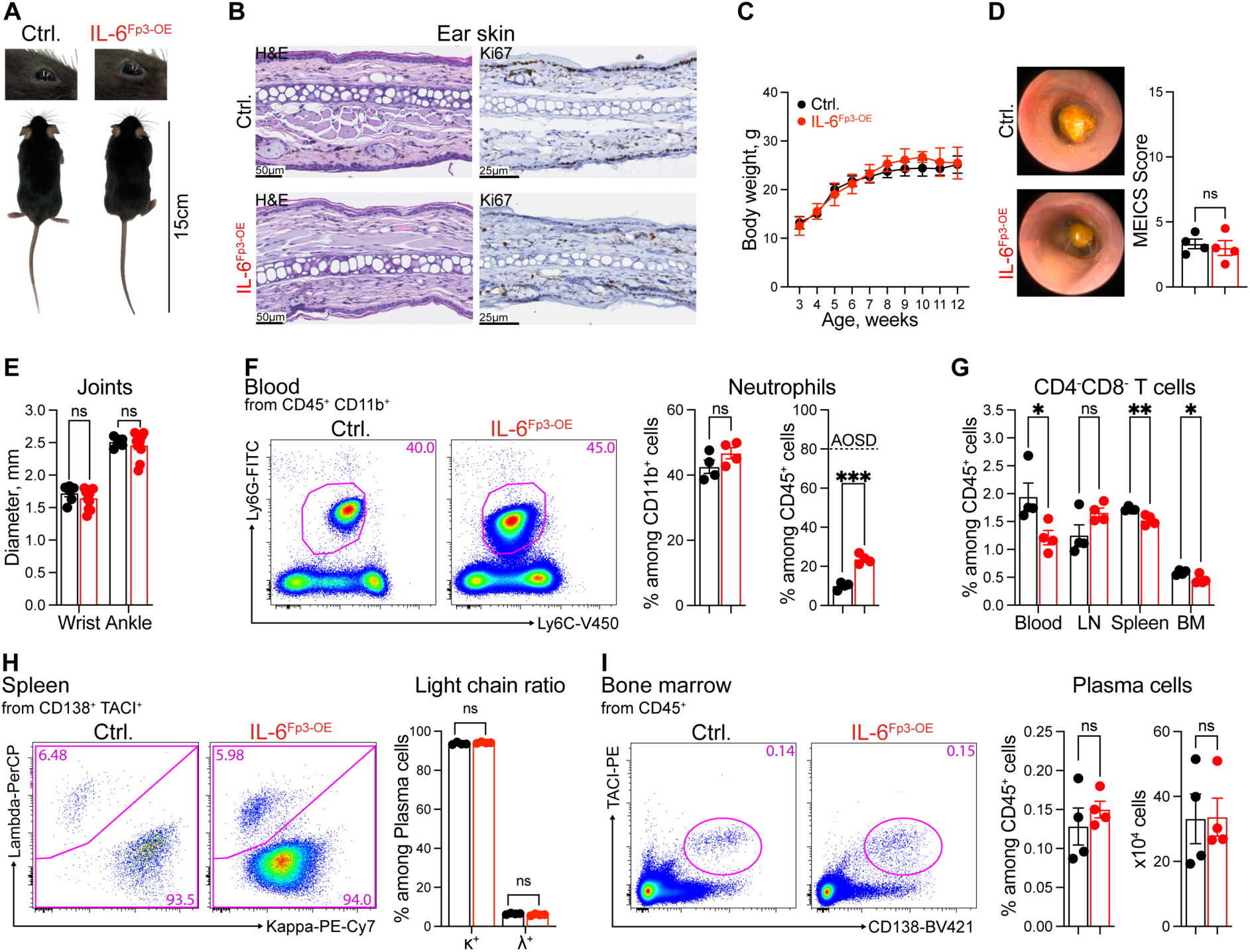
Analysis of exclusion diagnostic criteria for iMCD in IL-6^Fp3-OE^ mice. **A** – Macroscopic examination of 9 weeks old mice. **B** – Histological analysis (H&E and Ki67 staining) of the ear skin. Scale bars: 40 μm. n=4 mice per genotype, 8 weeks old. **C** – Body weight measurements at the indicated ages. n=10 mice per genotype. **D** – Representative pictures (video capture) from colonic mini-endoscopy and murine endoscopic index of colitis severity (MEICS) scores. n=4 mice per genotype, 8 weeks old. **E** – Thickness of the forepaw wrist and of the hind paw ankle of mice. n=6-11 mice per genotype, 9-10 weeks old. **F** – FACS analysis of Ly6C^int^Ly6G^hi^ neutrophils in the peripheral blood. (AOSD - adult-onset Still disease). n=4 mice per genotype, 9 weeks old. **G** – FACS analysis of CD4^-^CD8^-^ double negative T cells in the blood, lymph nodes (LN), spleen and bone marrow (BM). n=4 mice per genotype, 9 weeks old.**H** – FACS analysis of intracellular kappa (κ)- and lambda (λ)-light chain expression in plasma cells. n=4 mice per genotype, 8 weeks old. **I** – FACS analysis of CD138^+^TACI^+^ plasma cells in bone marrow of the left femur. n=4 mice per genotype, 9 weeks old. Graphs (D, E, F, G, H, I) show mean ± SEM. Each dot represents an individual mouse. P values were calculated using unpaired Student’s t-test and are indicated as follows: * (p < 0.05), ** (p < 0.01); ns – not significant (p > 0.05).

We next assessed autoimmune diseases that can mimic iMCD and must be excluded clinically^26^, including rheumatoid arthritis (RA), juvenile idiopathic arthritis (JIA), adult-onset Still disease (AOSD), and autoimmune lymphoproliferative syndrome (ALPS). Because RA and JIA are characterized by inflammatory joint swelling^45^, we measured wrist and ankle diameters and found no differences between IL-6^Fp3-OE^ and control mice (**Figure 4E, Supplemental Figure 4**).

Although Ly6C^int^ Ly6G^hi^ neutrophils were increased in IL-6^Fp3-OE^ mice, they accounted for less than 25% of circulating leukocytes (**Figure 4F**), well below the >80% required for AOSD^46^. Moreover, IL-6^Fp3-OE^ mice developed ANA autoantibodies (**Figure 3F**), whereas the Yamaguchi criteria require ANA negativity for AOSD diagnosis^47^. Likewise, CD4^-^CD8^-^ double-negative T cells were not enriched in the blood, lymph nodes, spleen, or bone marrow, excluding ALPS (**Figure 4G**).

Finally, we examined plasma cell neoplasms that can resemble iMCD, including multiple myeloma and polyneuropathy, organomegaly, endocrinopathy, monoclonal gammopathy, and skin changes (POEMS) syndrome. Both disorders are characterized by monoclonal plasma cell expansion^49,50^, whereas iMCD exhibits polyclonal plasmacytosis^26^. In non-neoplastic C57BL/6 mice, approximately 95% of plasma cells express κ light chains and 5% express λ light chains^51^. A substantial deviation from this distribution would indicate clonal expansion suggestive of malignancy. Previous models with transgenic IL-6 overexpression demonstrated that a C57BL/6 background prevents plasmacytoma development, resulting only in massive plasmacytosis^52^. Likewise, no light-chain restriction was observed in IL-6^Fp3-OE^ mice, indicating polyclonal plasmacytosis (**Figure 4H**). Furthermore, plasma cells were not increased in the bone marrow of IL-6^Fp3-OE^ mice (**Figure 4I**), whereas multiple myeloma is defined by ≥10% clonal plasma cells in this compartment^53^. The absence of λ light-chain restriction, together with the absence of detectable skin changes (**Figure 4A-B**), also argues against POEMS syndrome.

In summary, IL-6^Fp3-OE^ mice fulfilled the major and minor diagnostic criteria for iMCD while excluding relevant autoimmune disorders and plasma cell neoplasms^26^ and, therefore, represent a novel preclinical model for iMCD. Accordingly, IL-6^Fp3-OE^ mice are hereafter referred to as iMCD-like mice, and the two terms are used interchangeably throughout the manuscript.

### Plasma cell generation is dependent on classic IL-6 signaling in the context of iMCD-like disease

Having established the iMCD model, we next investigated the mechanisms underlying plasmacytosis in IL-6^Fp3-OE^ mice by examining IL-6 signaling through the IL-6Rα/gp130-STAT3 pathway^39^. Plasma cells from IL-6^Fp3-OE^ mice exhibited increased phosphorylated STAT3 (pSTAT3) mean fluorescence intensity (MFI) compared with controls, indicating active IL-6 signaling (**Figure 5A**).

**Figure 5.**
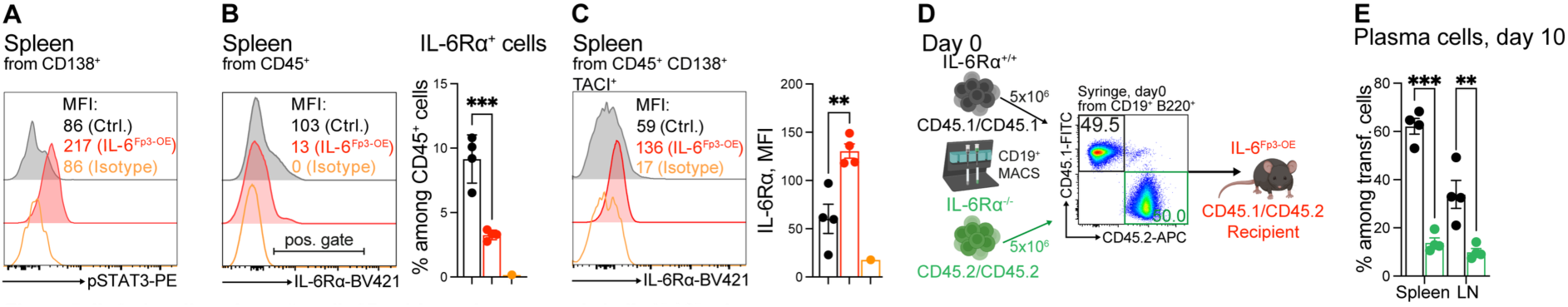
IL-6 signaling via surface IL-6Rα drives plasmacytosis in IL-6^Fp3-OE^ mice. **A** – FACS analysis of phosphorylated STAT3 (pSTAT3, Tyr705) levels in CD138^+^ plasma cells in the spleen. MFI - mean fluorescence intensity. **B** – FACS analysis of surface IL-6Rα expression on CD45^+^ leukocytes in the spleen. **C** – FACS analysis of surface IL-6Rα expression on CD138^+^TACI^+^ plasma cells in the spleen. **D** – Experimental set-up of CD19^+^ MACS-isolated IL-6Rα^+/+^ and IL-6Rα^-/-^ B cells co-transferred (1:1) into IL-6^Fp3-OE^ recipient mice. The FACS plot shows analysis of the mixed cell population in the syringe prior to injection. **E** – FACS analysis of plasma cell differentiation in the spleen and LN of IL-6^Fp3-OE^ recipient mice 10 days post-transfer. Graphs (B, C, E) show mean ± SEM. Each dot represents an individual mouse. P values were calculated using unpaired Student’s t-test (B, C, E) and are indicated as follows: *** (p < 0.05), ** (p < 0.01).

To determine the contribution of classic IL-6 signaling through membrane-bound IL-6Rα to plasmacytosis, we analyzed IL-6Rα expression on splenic leukocytes. Most immune cells in IL-6^Fp3-OE^ mice downregulated IL-6Rα expression, and only a small fraction of CD45⁺ cells retained surface IL-6Rα (**Figure 5B**), suggesting that the majority of leukocytes became unresponsive to classic IL-6 signaling. In contrast, plasma cells largely retained surface IL-6Rα expression, with higher IL-6Rα mean fluorescence intensity than control plasma cells (**Figure 5C**).

The combination of increased pSTAT3 and retained surface IL-6Rα expression indicates that plasma cells remain direct targets of classic IL-6 signaling during iMCD-like disease. To further assess the role of classic IL-6 signaling in plasma cell development, we co-transferred congenically distinct wild-type (IL-6Rα⁺^/^⁺) and IL-6Rα-deficient (IL-6Rα^-/-^) CD19⁺ B cells into IL-6^Fp3-OE^ recipient mice (**Figure 5D**). In this experimental setting, transferred IL-6Rα^-/-^ B cells specifically lack responsiveness to classic IL-6 signaling.

On the day of transfer, plasma cells were barely detectable in either donor preparation (**Supplemental Figure 5A**). Only minimal plasma cell differentiation occurred in healthy wild-type recipients (**Supplemental Figure 5B**). In contrast, a substantial fraction of IL-6Rα⁺^/^⁺ B cells differentiated into plasma cells in iMCD-like mice ten days after transfer (**Figure 5E**). Conversely, IL-6Rα^-/-^ B cells largely failed to generate plasma cells and were underrepresented in the spleen and lymph nodes of IL-6^Fp3-OE^ recipients compared with IL-6Rα⁺^/^⁺ donor-derived plasma cells (**Figure 5E**). These findings demonstrate that classic IL-6 signaling is required for efficient plasma cell differentiation during ongoing iMCD-like disease.

### Deficiency in soluble immunoglobulins protects IL-6^Fp3-OE^ mice from kidney dysfunction and significantly extends survival

Our findings suggest that IL-6^Fp3-OE^ mice develop lethal IgG1-driven plasmacytosis. To evaluate the impact of hypergammaglobulinemia on iMCD-like disease development and renal dysfunction, we used IgH^μγ1/μγ1^ mice, in which the μ constant region encodes only the membrane-bound form of IgM but not the secreted form^30^. Consequently, B cells in these mice are unable to undergo class switch recombination or secrete soluble immunoglobulins. We crossed IgH^μγ1/μγ1^ mice with IL-6^Fp3-OE^ mice to generate IgH^μγ1/μγ1^ IL-6^Fp3-OE^ mice, which overexpress IL-6 upon Foxp3-Cre-mediated transgene activation, similar to the original iMCD-like mice, but lack circulating immunoglobulins (**Figure 6A**). As expected, IgH^μγ1/μγ1^ IL-6^Fp3-OE^ mice maintained surface IgM expression but lacked IgD expression on B cells (**Supplemental Figure 6A**). Importantly, soluble IgM was absent from the serum of IgH^μγ1/μγ1^ IL-6^Fp3-OE^ mice (**Supplemental Figure 6B**). Similar to the original iMCD-like mice, IgH^μγ1/μγ1^ IL-6^Fp3-OE^ mice gained weight normally (**Supplemental Figure 6C**).

**Figure 6.**
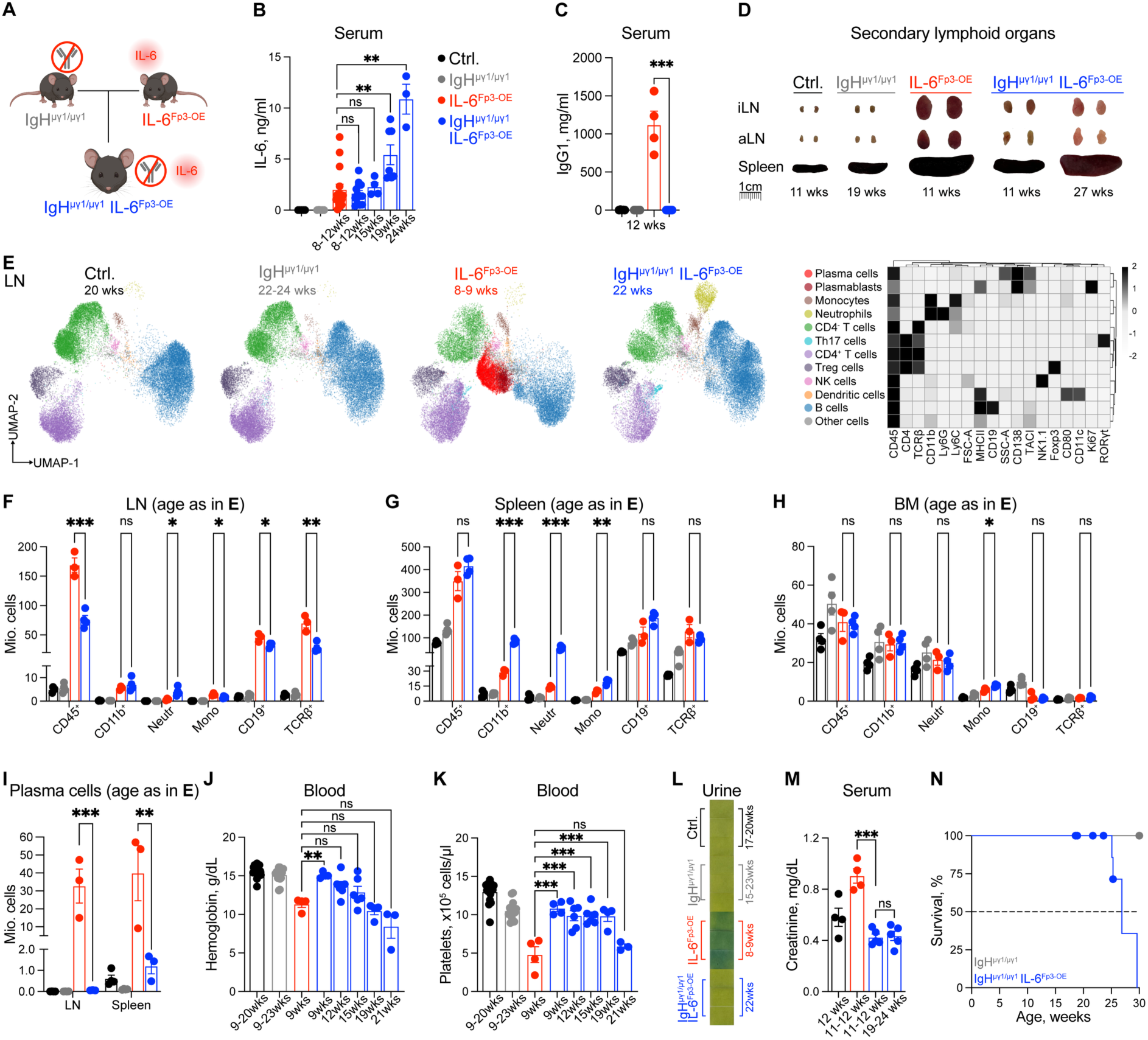
Soluble immunoglobulin-deficient IgH^μγ1/μγ1^ background protects IL-6^Fp3-OE^ mice from a plasmacytic iMCD-like disease. **A** – Generation of IL-6^Fp3-OE^ mice on the IgH^μγ1/μγ1^ background (IgH^μγ1/μγ1^ IL-6^Fp3-OE^ mice) with genetic inability to secrete soluble immunoglobulins. **B** – IL-6 levels in the serum of mice of the indicated genotypes and ages, measured by ELISA. **C** – IgG1 levels in the serum of 12-week-old mice of indicated genotypes, measured by ELISA. **D** – Macroscopic examination of inguinal lymph nodes (iLN), axillary lymph nodes (aLN), and spleens of mice of the indicated genotypes and ages. **E** – High-dimensional FACS analysis of immune cell populations (gated on live CD45^+^ cells) in lymph nodes (LN) of 20-week-old control mice, 22-24-week-old IgH^μγ1/μγ1^ mice, 8-9-week-old IL-6^Fp3-OE^, and 22-week-old IgH^μγ1/μγ1^ IL-6^Fp3-OE^ mice. The heat map displays respective individual clusters with median expression profiles identified by FlowSOM algorithm. A total of 30,000 cells per sample were used to generate UMAP. An overlay of the clusters identified by FlowSOM is shown. n=3 per group. **F** – FACS analysis of immune cell composition in lymph nodes of mice of the indicated genotypes and ages, as in **E**. Neutr. – Neutrophilic granulocytes (defined as CD45^+^ TCRβ^-^ CD19^-^ CD11b^+^ Ly6G^hi^ Ly6C^int^). Mono – Monocytes (defined as CD45^+^ TCRβ^-^ CD19^-^ CD11b^+^ Ly6G^neg^ Ly6C^int-hi^). **G** – FACS analysis of immune cell composition in the spleen of mice of the indicated genotypes and ages, as in **E**. Neutr. and mono., defined as in **F**. **H** – FACS analysis of immune cell composition of pooled bone marrow (two femora and one tibia) of mice of the indicated genotypes and age, as in **E**. Neutr. and mono., defined as in **F**. **I** – Number of CD138^+^TACI^+^ plasma cells in the lymph nodes and spleen of mice of the indicated genotypes and ages, as in **E**. **J** – Hemoglobin measurements of mice of the indicated genotypes and ages. **K** – Platelet counts of mice of the indicated genotypes and ages. **L** – Urine proteins of mice of the indicated genotypes and ages, measured by dipsticks (Roche Combur Test® HC). **M** – Creatinine levels in serum of mice of the indicated genotypes and ages, measured by a colorimetric assay. **N** – Kaplan-Meier survival curves of IgH^μγ1/μγ1^ (n=15) and IgH^μγ1/μγ1^ IL-6^Fp3-OE^ (n=17) mice. Inguinal, axial, and brachial lymph nodes were pooled and indicated as LN. Graphs (B, C, F, G, H, I, J, K, M) show mean ± SEM. Each dot represents an individual mouse. P values were calculated using unpaired Student’s t-test (C, F, G, H, I), one-way ANOVA (B, J, K, M), Gehan-Breslow-Wilcoxon-test (N), and are indicated as follows: * (p < 0.05), ** (p < 0.01), *** (p < 0.001); ns – not significant (p > 0.05).

To directly compare IL-6^Fp3-OE^ and IgH^μγ1/μγ1^ IL-6^Fp3-OE^ mice, we measured serum IL-6 levels and found similarly elevated levels in age-matched mice (**Figure 6B**), confirming that Foxp3-Cre-mediated IL-6 overexpression remained functional on the IgH^μγ1/μγ1^ background. IL-6 levels increased further with age in IgH^μγ1/μγ1^ IL-6^Fp3-OE^ mice and exceeded those observed in IL-6^Fp3-OE^ mice (**Figure 6B**). Nevertheless, as expected, IgH^μγ1/μγ1^ IL-6^Fp3-OE^ mice lacked serum IgG1 due to the IgH^μγ1/μγ1^ mutation (**Figure 6C**).

IL-6 is a potent mitogenic factor, and elevated IL-6 levels in IgH^μγ1/μγ1^ IL-6^Fp3-OE^ mice induced generalized lymphadenopathy, although to a lesser extent than in iMCD-like mice (**Figure 6D**). Splenomegaly progressed in adult animals, reaching the severity observed in iMCD-like mice (**Figure 6D, Supplemental Figure 6D**). To assess whether the enlargement of these organs in IgH^μγ1/μγ1^ IL-6^Fp3-OE^ mice resulted from expansion of specific immune cell populations, we analyzed the cellular composition of lymph nodes and spleens from 11-week- and 22-week-old IgH^μγ1/μγ1^ IL-6^Fp3-OE^ mice. Flow cytometry revealed that major immune cell populations were present in the secondary lymphoid organs of IgH^μγ1/μγ1^ IL-6^Fp3-OE^ mice (**Figure 6E, Supplemental Figure 6E**). Quantification of total cell numbers indicated that lymphadenopathy and splenomegaly arose from the outgrowth of CD19⁺ B cells and TCRβ⁺ T cells in IgH^μγ1/μγ1^ IL-6^Fp3-OE^ mice (**Figure 6F-G, Supplemental Figure 6F-G**). In addition, neutrophilia developed over time and was particularly pronounced in the spleens of 22-week-old IgH^μγ1/μγ1^ IL-6^Fp3-OE^ mice (**Figure 6G**). In contrast, bone marrow immune cell composition remained largely unchanged, irrespective of the IgH^μγ1/μγ1^ background (**Figure 6H**).

Notably, unlike IL-6^Fp3-OE^ mice, plasma cells in IgH^μγ1/μγ1^ IL-6^Fp3-OE^ mice represented only a minor population in secondary lymphoid organs and bone marrow (**Figure 6I, Supplemental Figure 6H-K**). This was unexpected, as the IgH^μγ1/μγ1^ mutation does not directly impair plasma cell development. To investigate this, we analyzed plasma cell maturation based on CD19 and B220 expression using a previously described gating strategy^54^. Nearly half of CD138⁺TACI⁺ cells in IgH^μγ1/μγ1^ IL-6^Fp3-OE^ mice displayed a plasmablast phenotype, compared with only 20% in control mice (**Supplemental Figure 7A**). Terminal plasma cell differentiation requires IRF4-mediated induction of Blimp-1 and subsequent repression of PAX^55,56^. However, CD138⁺TACI⁺ cells in IgH^μγ1/μγ1^ IL-6^Fp3-OE^ mice remained largely PAX5-positive, indicating incomplete differentiation (**Supplemental Figure 7B**). These data suggest that despite high levels of IL-6, IgH^μγ1/μγ1^ IL-6^Fp3-OE^ mice lack additional factor(s) required for sustained plasmacytosis.

The absence of plasmacytosis and hyperimmunoglobulinemia delayed anemia development in IgH^μγ1/μγ1^ IL-6^Fp3-OE^ mice (**Figure 6J**). Thrombocytopenia was also significantly reduced and reached levels observed in 9-week-old IL-6^Fp3-OE^ mice only by week 21 (**Figure 6K**). Importantly, IgH^μγ1/μγ1^ IL-6^Fp3-OE^ mice no longer developed proteinuria (**Figure 6L**), and serum creatinine levels remained normal (**Figure 6M**), indicating protection from renal dysfunction.

Despite high IL-6 levels, IgH^μγ1/μγ1^ IL-6^Fp3-OE^ mice did not succumb until 25 weeks of age (**Figure 6N**). Maximum survival was extended to 30 weeks, whereas none of the original iMCD-like mice with intact immunoglobulin secretion survived beyond 14 weeks. Because the major differences between IL-6^Fp3-OE^ and IgH^μγ1/μγ1^ IL-6^Fp3-OE^ mice are the inability to secrete soluble immunoglobulins and the absence of severe plasmacytosis in the latter strain, we conclude that IL-6-driven IgG1 plasmacytosis is a key determinant of lethal disease in iMCD-like mice.

## Discussion

iMCD is a rare, life-threatening lymphoproliferative disorder with an estimated annual incidence and prevalence of 3.4 and 6.9 cases per million individuals in the USA, respectively. The mechanisms underlying iMCD remain unknown, and treatment options are limited. The only approved, albeit non-curative, therapy is IL-6 blockade^57^. IL-6 is a key driver of plasma cell development, and the best responses to anti-IL-6 therapy are observed in the plasmacytic (iMCD-IPL) subtype. However, treatment requires lifelong administration negatively affecting patients’ quality of life even among good responders^40^. A better understanding of iMCD pathogenesis is essential for developing novel therapies, for which preclinical models are invaluable.

Previous attempts to develop IL-6-based iMCD models were limited to transgenic mice with constitutive overexpression of IL-6. Although some models reproduced selected features of iMCD^52,58–61^, others developed primary morbidities unrelated to the disease^62–72^. Moreover, these models are difficult to maintain as breeding colonies due to their severe phenotype and shortened lifespan. To our knowledge, no IL-6-based mouse model has been evaluated against the most recent international consensus diagnostic criteria for iMCD^26^. Here, we developed IL-6^Fp3-OE^ mice with conditional IL-6 overexpression in Foxp3^+^ Treg cells, which faithfully recapitulate clinical, hematologic, and histopathological features of human iMCD.

The cellular source of pathogenic IL-6 in iMCD remains unknown. B cells within germinal centers have been proposed as one source^73^, whereas later studies identified plasma cells as the predominant IL-6-expressing cells in iMCD-IPL and vascular endothelial cells in iMCD-TAFRO^74^. However, IL-6 is a soluble cytokine that can induce its own expression through paracrine signaling^36^, making it difficult to identify the initiating cellular source. We selected Treg cells as IL-6-producing cells to model iMCD because immunohistochemical analyses of lymph node biopsies from iMCD patients revealed an abundance of FOXP3-positive cells adjacent to plasma cells, the key pathogenic cell type in iMCD-IPL. The enrichment of FOXP3-positive cells near plasma cells raises the possibility that germinal center Treg cells normally restrain excessive plasma cell responses within this microenvironment. However, our patient cohort was too small to draw definitive conclusions regarding Treg cell abundance or function across iMCD subtypes, and future studies will be required to address this question. Nevertheless, targeting IL-6 overexpression to Treg cells generated the IL-6^Fp3-OE^ mouse model, which fulfilled both major and five of the 11 minor consensus diagnostic criteria for iMCD with complete penetrance. Moreover, the full iMCD-like phenotype developed by 8 weeks of age, compared with 15–18 weeks in previous models, which also exhibited incomplete penetrance^60,61^. Importantly, our model does not suggest that Treg cells are the physiological source of IL-6 in human disease. Rather, it demonstrates that sustained IL-6 production within a plasma cell-rich microenvironment is sufficient to drive an iMCD-like syndrome.

The relationship between iMCD and HHV-8-associated multicentric Castleman disease is of particular interest because both conditions share elevated IL-6 signaling, plasmacytosis, and systemic inflammation^26^. HHV-8 encodes a viral IL-6 homolog capable of activating many of the same downstream pathways as human IL-6. Recently, treatment with CD19 CAR T cells successfully eliminated IL-6-producing B cells in a patient with iMCD-IPL, leading to sustained remission^75^. Our findings further support persistent IL-6 activity as a common pathogenic mechanism underlying Castleman disease pathology.

Clinically, IL-6^Fp3-OE^ mice recapitulate key hallmarks of iMCD-IPL, including plasmacytosis, hypergammaglobulinemia, and anemia. They also exhibit features overlapping with iMCD-TAFRO syndrome, such as thrombocytopenia, hypervascularization, renal dysfunction, and organomegaly. Unlike iMCD-TAFRO, which typically presents with acute onset and lacks plasmacytic histopathology^76^, IL-6^Fp3^ ^OE^ mice develop a chronic, progressive disease accompanied by marked plasmacytosis.

In our model, plasmacytosis is driven by IL-6 acting on plasma cells through classic IL-6 signaling via membrane-bound IL-6Rα, leading to STAT3 phosphorylation. This plasmacytosis results in profound IgG1 hypergammaglobulinemia, promoting immune complex deposition in the kidneys and consequent renal dysfunction. Accordingly, IL-6^Fp3-OE^ mice exhibit both structural renal pathology and functional impairment. Consistent with these findings, previous reports showed that IL-6 transgenic mice with high IgG1 levels developed mesangio-proliferative glomerulonephritis^58,59^.

To directly test the pathogenic role of hypergammaglobulinemia, we crossed IL-6^Fp3-OE^ mice onto an IgH^μγ1/μγ1^ background, thereby genetically blocking B cell class-switching and the secretion of soluble immunoglobulins. The resulting IgH^μγ1/μγ1^ IL-6^Fp3-OE^ mice displayed high levels of IL-6 and undetectable serum immunoglobulins. Unexpectedly, however, they also failed to develop plasmacytosis, suggesting that soluble immunoglobulins contribute to plasma cell development^77^. The IgH^μγ1/μγ1^ background resulted in less pronounced lymphadenopathy and delayed onset of splenomegaly. Interestingly, splenic enlargement in IgH^μγ1/μγ1^ IL-6^Fp3-OE^ mice was driven by increased numbers of T cells, B cells, and neutrophils. This suggests that, in the absence of plasmacytosis, chronic IL-6 promotes alternative inflammatory cell expansion, including neutrophilia. Anemia and thrombocytopenia were significantly delayed, suggesting that autoantibodies in the original IL-6^Fp3-OE^ mice may promote abnormal hematologic parameters.

A notable finding of our study was that the IgH^μγ1/μγ1^ background completely rescued proteinuria and normalized serum creatinine levels. As we did not observe plasmacytosis in the kidneys of the original IL-6^Fp3-OE^ mice, this supports the pathogenic role of immune complexes in renal involvement. However, our data do not suggest that immunoglobulins are the sole drivers of organ damage in iMCD. Despite marked protection in mice unable to secrete antibodies, chronic IL-6 exposure ultimately remained detrimental. Thus, hypergammaglobulinemia, immune complex deposition, and direct IL-6-mediated inflammatory effects are all likely to contribute to disease pathology.

In summary, our study identifies aberrant IL-6-driven IgG1 plasmacytosis as a key pathogenic mechanism in iMCD-like disease and provides a robust preclinical model for investigating iMCD pathogenesis and evaluating therapeutic interventions beyond IL-6 blockade.

## Acknowledgements

We thank Dr. Tobias Bopp for providing Foxp3-Cre mice on a C57BL/6 background (originally from Dr. Shimon Sakaguchi), Dr. Carsten Deppermann for fruitful discussions on complete blood count analysis, and Dr. Nadine Hövelmeyer for valuable discussions on B-cell biology. We also thank Dr. Khalad Karram for assistance with fluorescence microscopy, Elena Zurkowski for excellent technical assistance with the evaluation of gut inflammation, and Claudia Braun and Bonny Adami for preparing histological sections and performing stainings (all from the University Medical Center Mainz). I.A.M. is personally grateful to the members of the Bluestone laboratory (Dr. Jeffrey Bluestone, Dr. James Lee, and Dr. Caroline Raffin (Mufazalov)) at the Diabetes Center, University of California, San Francisco (UCSF), for their support during the initiation of this project.

This work was supported by the Deutsche Forschungsgemeinschaft (DFG, German Research Foundation) through grants 318346496 – SFB 1292 and 490846870 – TRR355 to A.W. M.M.G. and A.H. were supported through DFG grant 318346496 – SFB 1292. T.K. was supported by the DFG (TRR274 (ID 408885537), TRR355 (ID 490846870), GRK2668 (ID 435874434), and EXC 2145 (SyNergy, ID 390857198)), by the European Research Council (ERC-ADG “Breaking Bad”), and by the Hertie Network of Clinical Neuroscience. Additional funding was provided by the interdisciplinary life-science research initiative ReALity (Resilience, Adaptation and Longevity) of the Johannes Gutenberg University Mainz to support I.A.M.

We thank the FACS Core Facility CFFC PKZI FZI of the University Medical Center Mainz for providing access to the FACSymphony A5, funded by the DFG (INST 371/45-1 FUGB).

Experimental design schemes were created with BioRender.com.

## Authorship Contributions

D.C.U. designed and performed the experiments, analyzed and interpreted data, and wrote the manuscript; D.A., L.R.A., H.S.S., M.H., M.B., C.S., Z.E., A.P.Z., G.A.B., and D.C.N. performed experiments; A.H., S.Z., and M.M.G. performed and analyzed histological data; S.U-S. and K.J. performed laboratory blood analyses; F.T.W and J.C.A-F. contributed essential ideas and reagents; T.K. contributed essential ideas and revised the manuscript; A.W. contributed essential ideas, advised on experiments, provided mice, and provided funding support; D.A., F.T.W and A.W. critically edited the manuscript; I.A.M. supervised the study, designed experiments, interpreted data and wrote the manuscript.

## References

1. Yu L, Tu M, Cortes J, et al. Clinical and pathological characteristics of HIV- and HHV-8-negative Castleman disease. Blood. 2017;129(12):1658–1668.

2. Liu AY, Nabel CS, Finkelman BS, et al. Idiopathic multicentric Castleman’s disease: a systematic literature review. Lancet Haematol. 2016;3(4):e163–175.

3. Dispenzieri A, Fajgenbaum DC. Overview of Castleman disease. Blood. 2020;135(16):1353–1364.

4. Nishimoto N, Kanakura Y, Aozasa K, et al. Humanized anti-interleukin-6 receptor antibody treatment of multicentric Castleman disease. Blood. 2005;106(8):2627–2632.

5. van Rhee F, Casper C, Voorhees PM, et al. A phase 2, open-label, multicenter study of the long-term safety of siltuximab (an anti-interleukin-6 monoclonal antibody) in patients with multicentric Castleman disease. Oncotarget. 2015;6(30):30408–30419.

6. Yin X, Liu Y, Lv Z, et al. scRNA-seq reveals the landscape of immune repertoire of PBMNCs in iMCD. Oncogene. 2024;43(37):2795–2805.

7. Harada T, Kikushige Y, Miyamoto T, et al. Peripheral helper-T-cell-derived CXCL13 is a crucial pathogenic factor in idiopathic multicentric Castleman disease. Nat Commun. 2023;14(1):6959.

8. Pierson SK, Katz L, Williams R, et al. CXCL13 is a predictive biomarker in idiopathic multicentric Castleman disease. Nat Commun. 2022;13(1):7236.

9. Pierson SK, Stonestrom AJ, Shilling D, et al. Plasma proteomics identifies a ‘chemokine storm’ in idiopathic multicentric Castleman disease. Am J Hematol. 2018;93(7):902–912.

10. Srkalovic G, Nijim S, Srkalovic MB, Fajgenbaum D. Increase in Vascular Endothelial Growth Factor (VEGF) Expression and the Pathogenesis of iMCD-TAFRO. Biomedicines. 2024;12(6).

11. Mumau MD, Gonzalez MV, Ma C, et al. Identifying and Targeting TNF Signaling in Idiopathic Multicentric Castleman’s Disease. N Engl J Med. 2025;392(6):616–618.

12. Endo Y, Koga T, Umeda M, Furukawa K, Takenaka M, Kawakami A. Successful canakinumab treatment for activated innate response in idiopathic Castleman’s disease with multiple heterozygous MEFV exon 2 variants. Clin Immunol. 2020;219:108547.

13. Soudet S, Fajgenbaum D, Delattre C, et al. Schnitzler syndrome co-occurring with idiopathic multicentric Castleman disease that responds to anti-IL-1 therapy: A case report and clue to pathophysiology. Curr Res Transl Med. 2018;66(3):83–86.

14. El-Osta H, Janku F, Kurzrock R. Successful treatment of Castleman’s disease with interleukin-1 receptor antagonist (Anakinra). Mol Cancer Ther. 2010;9(6):1485–1488.

15. Galeotti C, Tran TA, Franchi-Abella S, Fabre M, Pariente D, Kone-Paut I. IL-1RA agonist (anakinra) in the treatment of multifocal castleman disease: case report. J Pediatr Hematol Oncol. 2008;30(12):920–924.

16. Sultan AW, Griesshammer E, Hinrichs C, et al. TAFRO syndrome requiring combined IL 6 and IL 1 inhibition: a case report. Front Immunol. 2025;16:1729525.

17. Palmeri S, Ferro J, Natoli V, et al. Efficacy of High-Dose Intravenous Anakinra in Pediatric TAFRO Syndrome: Report of Two Cases and Literature Review. Pediatr Blood Cancer. 2025;72(8):e31759.

18. Liu YT, Gao YH, Zhao H, et al. Sirolimus is effective for refractory/relapsed idiopathic multicentric Castleman disease: A single-center, retrospective study. Ann Hematol. 2024;103(10):4223–4230.

19. Fajgenbaum DC, Langan RA, Japp AS, et al. Identifying and targeting pathogenic PI3K/AKT/mTOR signaling in IL-6-blockade-refractory idiopathic multicentric Castleman disease. J Clin Invest. 2019;129(10):4451–4463.

20. Koga T, Sumiyoshi R, Shimizu T, et al. A Placebo-Controlled Exploratory Trial of Sirolimus for Tocilizumab-Resistant Idiopathic Multicentric Castleman Disease: Early Termination and Long-Term Extension Results Based on Descriptive Results From Two Patients. Cureus. 2025;17(12):e98233.

21. Zhao H, Zhang MY, Shen KN, et al. A phase 2 prospective study of bortezomib, cyclophosphamide, and dexamethasone in patients with newly diagnosed iMCD. Blood. 2023;141(21):2654–2657.

22. Kakutani T, Nunokawa T, Chinen N, Tamai Y. Treatment-resistant idiopathic multicentric Castleman disease with thrombocytopenia, anasarca, fever, reticulin fibrosis, renal dysfunction, and organomegaly managed with Janus kinase inhibitors: A case report. Medicine (Baltimore). 2022;101(48):e32200.

23. Gao YH, Li SY, Dang Y, Duan MH, Zhang L, Li J. Efficacy and safety of orelabrutinib in relapsed/refractory idiopathic multicentric Castleman disease: A single-centre, retrospective study. Br J Haematol. 2025;206(1):152–158.

24. Fajgenbaum DC. Novel insights and therapeutic approaches in idiopathic multicentric Castleman disease. Blood. 2018;132(22):2323–2330.

25. Pierson SK, Khor JS, Ziglar J, et al. ACCELERATE: A Patient-Powered Natural History Study Design Enabling Clinical and Therapeutic Discoveries in a Rare Disorder. Cell Rep Med. 2020;1(9):100158.

26. Fajgenbaum DC, Uldrick TS, Bagg A, et al. International, evidence-based consensus diagnostic criteria for HHV-8-negative/idiopathic multicentric Castleman disease. Blood. 2017;129(12):1646–1657.

27. Mufazalov IA, Andruszewski D, Schelmbauer C, et al. Cutting Edge: IL-6-Driven Immune Dysregulation Is Strictly Dependent on IL-6R alpha-Chain Expression. J Immunol. 2020;204(4):747–751.

28. Knopp T, Jung R, Wild J, et al. Myeloid cell-derived interleukin-6 induces vascular dysfunction and vascular and systemic inflammation. Eur Heart J Open. 2024;4(4):oeae046.

29. Wing K, Onishi Y, Prieto-Martin P, et al. CTLA-4 control over Foxp3+ regulatory T cell function. Science. 2008;322(5899):271–275.

30. Waisman A, Kraus M, Seagal J, et al. IgG1 B cell receptor signaling is inhibited by CD22 and promotes the development of B cells whose survival is less dependent on Ig alpha/beta. J Exp Med. 2007;204(4):747–758.

31. Wunderlich FT, Strohle P, Konner AC, et al. Interleukin-6 signaling in liver-parenchymal cells suppresses hepatic inflammation and improves systemic insulin action. Cell Metab. 2010;12(3):237–249.

32. Caton ML, Smith-Raska MR, Reizis B. Notch-RBP-J signaling controls the homeostasis of CD8-dendritic cells in the spleen. J Exp Med. 2007;204(7):1653–1664.

33. Reissig S, Tang Y, Nikolaev A, et al. Elevated levels of Bcl-3 inhibits Treg development and function resulting in spontaneous colitis. Nat Commun. 2017;8:15069.

34. Shanmugavadivu A, Carter K, Zonouzi AP, Waisman A, Regen T. Protocol for the collection and analysis of the different immune cell subsets in the murine intestinal lamina propria. STAR Protoc. 2024;5(3):103154.

35. Schaffer S, Maul-Pavicic A, Voll RE, Chevalier N. Optimized isolation of renal plasma cells for flow cytometric analysis. J Immunol Methods. 2019;474:112628.

36. Grebenciucova E, VanHaerents S. Interleukin 6: at the interface of human health and disease. Front Immunol. 2023;14:1255533.

37. Pierson SK, Brandstadter JD, Torigian DA, et al. Characterizing the heterogeneity of Castleman disease and oligocentric subtype: findings from the ACCELERATE registry. Blood Adv. 2025;9(8):1952–1965.

38. Glatman Zaretsky A, Konradt C, Depis F, et al. T Regulatory Cells Support Plasma Cell Populations in the Bone Marrow. Cell Rep. 2017;18(8):1906–1916.

39. Garbers C, Heink S, Korn T, Rose-John S. Interleukin-6: designing specific therapeutics for a complex cytokine. Nat Rev Drug Discov. 2018;17(6):395–412.

40. Bustamante MS, Pierson SK, Ren Y, et al. Longitudinal, natural history study reveals the disease burden of idiopathic multicentric Castleman disease. Haematologica. 2024;109(7):2196–2206.

41. Sawada E, Shioda Y, Ogawa K, et al. A Case of Castleman’s Disease with a Marked Infiltration of IgG4-Positive Cells in the Renal Interstitium. Diagnostics (Basel). 2024;14(5).

42. Kawanishi M, Kamei F, Sonoda H, et al. Utility of renal biopsy in differentiating idiopathic multicentric Castleman disease from IgG4-related disease. CEN Case Rep. 2023;12(2):242–248.

43. Sakaguchi S, Sakaguchi N, Asano M, Itoh M, Toda M. Immunologic self-tolerance maintained by activated T cells expressing IL-2 receptor alpha-chains (CD25). Breakdown of a single mechanism of self-tolerance causes various autoimmune diseases. J Immunol. 1995;155(3):1151–1164.

44. Godfrey VL, Wilkinson JE, Rinchik EM, Russell LB. Fatal lymphoreticular disease in the scurfy (sf) mouse requires T cells that mature in a sf thymic environment: potential model for thymic education. Proc Natl Acad Sci U S A. 1991;88(13):5528–5532.

45. Martini A, Lovell DJ, Albani S, et al. Juvenile idiopathic arthritis. Nat Rev Dis Primers. 2022;8(1):5.

46. Tomaras S, Goetzke CC, Kallinich T, Feist E. Adult-Onset Still’s Disease: Clinical Aspects and Therapeutic Approach. J Clin Med. 2021;10(4).

47. Yamaguchi M, Ohta A, Tsunematsu T, et al. Preliminary criteria for classification of adult Still’s disease. J Rheumatol. 1992;19(3):424–430.

48. Rieux-Laucat F, Magerus-Chatinet A. Autoimmune lymphoproliferative syndrome: a multifactorial disorder. Haematologica. 2010;95(11):1805–1807.

49. Dispenzieri A. POEMS Syndrome: 2019 Update on diagnosis, risk-stratification, and management. Am J Hematol. 2019;94(7):812–827.

50. Malard F, Neri P, Bahlis NJ, et al. Multiple myeloma. Nature Reviews Disease Primers. 2024;10(1):45.

51. Haughton G, Lanier LL, Babcock GF. The murine kappa light chain shift. Nature. 1978;275(5676):154–157.

52. Suematsu S, Matsusaka T, Matsuda T, et al. Generation of plasmacytomas with the chromosomal translocation t(12;15) in interleukin 6 transgenic mice. Proc Natl Acad Sci U S A. 1992;89(1):232–235.

53. Rajkumar SV. Multiple myeloma: 2022 update on diagnosis, risk stratification, and management. Am J Hematol. 2022;97(8):1086–1107.

54. Pracht K, Meinzinger J, Daum P, et al. A new staining protocol for detection of murine antibody-secreting plasma cell subsets by flow cytometry. Eur J Immunol. 2017;47(8):1389–1392.

55. Klein U, Casola S, Cattoretti G, et al. Transcription factor IRF4 controls plasma cell differentiation and class-switch recombination. Nature Immunology. 2006;7(7):773–782.

56. Nutt SL, Hodgkin PD, Tarlinton DM, Corcoran LM. The generation of antibody-secreting plasma cells. Nature Reviews Immunology. 2015;15(3):160–171.

57. Mukherjee S, Martin R, Sande B, Paige JS, Fajgenbaum DC. Epidemiology and treatment patterns of idiopathic multicentric Castleman disease in the era of IL-6-directed therapy. Blood Adv. 2022;6(2):359–367.

58. Suematsu S, Matsuda T, Aozasa K, et al. IgG1 plasmacytosis in interleukin 6 transgenic mice. Proc Natl Acad Sci U S A. 1989;86(19):7547–7551.

59. Brandt SJ, Bodine DM, Dunbar CE, Nienhuis AW. Dysregulated interleukin 6 expression produces a syndrome resembling Castleman’s disease in mice. J Clin Invest. 1990;86(2):592–599.

60. Katsume A, Saito H, Yamada Y, et al. Anti-interleukin 6 (IL-6) receptor antibody suppresses Castleman’s disease like symptoms emerged in IL-6 transgenic mice. Cytokine. 2002;20(6):304–311.

61. Suthaus J, Stuhlmann-Laeisz C, Tompkins VS, et al. HHV-8-encoded viral IL-6 collaborates with mouse IL-6 in the development of multicentric Castleman disease in mice. Blood. 2012;119(22):5173–5181.

62. Turksen K, Kupper T, Degenstein L, Williams I, Fuchs E. Interleukin 6: insights to its function in skin by overexpression in transgenic mice. Proc Natl Acad Sci U S A. 1992;89(11):5068–5072.

63. Woodroofe C, Muller W, Ruther U. Long-term consequences of interleukin-6 overexpression in transgenic mice. DNA Cell Biol. 1992;11(8):587–592.

64. Campbell IL, Abraham CR, Masliah E, et al. Neurologic disease induced in transgenic mice by cerebral overexpression of interleukin 6. Proc Natl Acad Sci U S A. 1993;90(21):10061–10065.

65. Campbell IL, Hobbs MV, Dockter J, Oldstone MB, Allison J. Islet inflammation and hyperplasia induced by the pancreatic islet-specific overexpression of interleukin-6 in transgenic mice. Am J Pathol. 1994;145(1):157–166.

66. DiCosmo BF, Geba GP, Picarella D, et al. Airway epithelial cell expression of interleukin-6 in transgenic mice. Uncoupling of airway inflammation and bronchial hyperreactivity. J Clin Invest. 1994;94(5):2028–2035.

67. Fattori E, Lazzaro D, Musiani P, Modesti A, Alonzi T, Ciliberto G. IL-6 expression in neurons of transgenic mice causes reactive astrocytosis and increase in ramified microglial cells but no neuronal damage. Eur J Neurosci. 1995;7(12):2441–2449.

68. Kitamura H, Kawata H, Takahashi F, Higuchi Y, Furuichi T, Ohkawa H. Bone marrow neutrophilia and suppressed bone turnover in human interleukin-6 transgenic mice. A cellular relationship among hematopoietic cells, osteoblasts, and osteoclasts mediated by stromal cells in bone marrow. Am J Pathol. 1995;147(6):1682–1692.

69. Tsujinaka T, Ebisui C, Fujita J, et al. Muscle undergoes atrophy in association with increase of lysosomal cathepsin activity in interleukin-6 transgenic mouse. Biochem Biophys Res Commun. 1995;207(1):168–174.

70. De Benedetti F, Alonzi T, Moretta A, et al. Interleukin 6 causes growth impairment in transgenic mice through a decrease in insulin-like growth factor-I. A model for stunted growth in children with chronic inflammation. J Clin Invest. 1997;99(4):643–650.

71. Gorshkova EA, Zvartsev RV, Drutskaya MS, Gubernatorova EO. [Humanized Mouse Models as a Tool to Study Proinflammatory Cytokine Overexpression]. Mol Biol (Mosk). 2019;53(5):755–773.

72. Jergovic M, Thompson HL, Bradshaw CM, et al. IL-6 can singlehandedly drive many features of frailty in mice. Geroscience. 2021;43(2):539–549.

73. Yoshizaki K, Matsuda T, Nishimoto N, et al. Pathogenic significance of interleukin-6 (IL-6/BSF-2) in Castleman’s disease. Blood. 1989;74(4):1360–1367.

74. Nishikori A, Nishimura MF, Nishimura Y, et al. Distinct interleukin-6 production in IPL and TAFRO subtypes of idiopathic multicentric Castleman disease. Haematologica. 2026;111(5):1705–1715.

75. Zhang Y, Cheng F, Deng B, et al. Sustained Complete Response in Refractory Idiopathic Multicentric Castleman Disease Following a Single CD19 CAR T-Cell Infusion: A Case Report With Mechanistic Insight. Hematol Oncol. 2026;44(4):e70214.

76. Nishimura Y, Fajgenbaum DC, Pierson SK, et al. Validated international definition of the thrombocytopenia, anasarca, fever, reticulin fibrosis, renal insufficiency, and organomegaly clinical subtype (TAFRO) of idiopathic multicentric Castleman disease. Am J Hematol. 2021;96(10):1241–1252.

77. Nguyen TTT, Graf BA, Randall TD, Baumgarth N. sIgM-FcmuR Interactions Regulate Early B Cell Activation and Plasma Cell Development after Influenza Virus Infection. J Immunol. 2017;199(5):1635–1646.

