## Supplemental Materials for "Targeting soluble immunoglobulins ameliorates idiopathic multicentric Castleman disease in a mouse model"

**Table of Contents:**

Supplemental Methods
Supplemental Table 1
Supplemental Figure 1-7
Supplemental References

### Supplemental Methods

#### Mice breeding strategies

Mice carrying a murine *Il6* cDNA transgene with a STOP cassette inserted into the *Rosa26* locus ($Rosa26^{IL\text{-}6(stop/stop)}$ mice) and designed for Cre-mediated activation of irreversible IL-6 overexpression ($IL\text{-}6^{OE}$) were previously generated by us[1,2]. These $Rosa26^{IL\text{-}6(stop/stop)}$ mice were crossed with Foxp3-Cre mice[3], and subsequent intercrosses produced male offspring homozygous for the *Il6* transgene and hemizygous for Foxp3-Cre ($IL\text{-}6^{Fp3\text{-}OE}$) which were used as experimental animals. Their male littermates lacking Foxp3-Cre were used as controls (Ctrl.) (**Supplemental Figure 1**).

To generate IL-6-overexpressing mice unable to secrete soluble immunoglobulins, $IL\text{-}6^{Fp3\text{-}OE}$ mice were crossed with mice carrying a mutated IgH locus ($IgH^{\mu\gamma1/\mu\gamma1}$ mice[4]), resulting in the generation of $IgH^{\mu\gamma1/\mu\gamma1}$ $IL\text{-}6^{Fp3\text{-}OE}$ mice. A breeding strategy similar to that used for $IL\text{-}6^{Fp3\text{-}OE}$ mice was employed to maintain the colony and generate experimental $IgH^{\mu\gamma1/\mu\gamma1}$ $IL\text{-}6^{Fp3\text{-}OE}$ mice and their littermate $IgH^{\mu\gamma1/\mu\gamma1}$ control mice.

IL-6Rα full-knockout ($IL\text{-}6R\alpha^{-/-}$) mice were selected from the breeding between *Il6ra* flox mice[5] and CD11c-Cre mice[6], which displayed occasional spontaneous germline Cre activity in our previous study[1].

All mice were maintained on a C57BL/6 background and bred in-house under specific pathogen-free (SPF) conditions at the Translational Animal Research Center (TARC), University Medical Center of Johannes Gutenberg University Mainz.

#### ELISA and creatinine measurement in murine samples

IL-6 levels in serum and tissues were measured using the BD OptEIA™ Mouse IL-6 ELISA Set according to the manufacturer's instructions. Spleen and inguinal lymph node samples were lysed in RIPA buffer (Thermo Scientific™) and stored at -80°C prior to analysis.

Serum IgG1, IgE, and IgM levels were determined by ELISA using in-house standards. Briefly, 96-well plates were coated with goat anti-mouse IgG1 (Cat. No. 1070-01), rat anti-mouse IgE (Cat. No. 1130-01), or goat anti-mouse IgM (Cat. No. 1020-01) antibodies (all from SouthernBiotech). After blocking with BSA, serially diluted serum samples were added together with biotinylated rat anti-mouse detection antibodies specific for IgG1 (Cat. No. 1144-08), IgE (Cat. No. 1130-08), or IgM (Cat. No. 1140-08) (all from SouthernBiotech). Detection was performed using streptavidin-conjugated alkaline phosphatase (Roche, Cat. No. 11089161001) and p-nitrophenyl phosphate substrate (Sigma-Aldrich, Cat. No. 487666-1EA).

Serum creatinine levels were measured using the Creatinine Assay Kit (Creative Biomart, Kit-0261) according to the manufacturer's instructions. Optical density was measured using an Infinite M200 PRO NanoQuant reader (Tecan).

#### Flow cytometry with murine cells

Single-cell suspensions from lymphoid and non-lymphoid organs in phosphate-buffered saline (PBS) supplemented with 2% fetal calf serum (FCS) were kept on ice. Erythrocytes were lysed using ACK buffer (50 mM ammonium chloride, 10 mM potassium bicarbonate, and 1 mM EDTA) for 3-5 minutes at room temperature. For FACS analysis, Fc receptors were first blocked using Fc-Block (5 µg/mL; BioXcell) for 15 minutes at 4 °C. Next, the fixable viability dye FVS-UV440 (BD) was applied in PBS for 15 minutes prior to antibody staining. When FVS-eFluor780 or FVS-eFluor506 (eBioscience) were used, the viability dye was included together with the antibody staining step. Cells were then stained with fluorochrome- or biotin-conjugated antibodies in PBS supplemented with 2% FCS for 20 minutes.

For intracellular staining of transcription factors, the Foxp3/Transcription Factor Staining Buffer Set (eBioscience) was used according to the manufacturer's recommendations. In short, after fixation, intracellular antibodies and fluorochrome-conjugated streptavidin diluted in permeabilization buffer were incubated with cells at 4 °C for 1 hour or overnight.

To simultaneously detect GFP and transcription factors, cells were first stained with anti-TCRβ and anti-CD4 surface antibodies, then fixed and permeabilized using the Cytofix/Cytoperm kit (BD Biosciences), followed by staining with anti-GFP antibodies. The cells were then further fixed and permeabilized using the Foxp3/Transcription Factor Staining Buffer Set (eBioscience) and stained for Foxp3. FACS antibodies used in this study are summarized in Supplemental Table 1.

Samples were acquired on a BD FACSCanto™ II, BD FACSymphony™ A5, or BD FACSymphony™ A5 SE, and analyzed using FlowJo™ 10.8.1 software. Doublets were excluded based on FSC and SSC properties, and dead cells were excluded by gating on fixable viability dye–negative cells.

For high-dimensional flow cytometry analysis, appropriate pre-gating was applied and equally downsampled FCS files were merged. The FlowSOM algorithm was used to identify meta-clusters[7], and UMAP-clustering was performed for dimensionality reduction and visualization[8]. Subsequently, data were exported from FlowJo as CSV files and processed in the R environment (R Studio version 025.09.0+387) for visualization and layout adjustments using the ggplot2 and pheatmap packages[9,10].

**Murine B cell isolation and transfer**

Spleens were harvested from naïve wild type IL-6Rα$^{+/+}$ and IL-6Rα$^{-/-}$ donor mice on CD45.1/CD45.1 and CD45.2/CD45.2 background, respectively. Single-cell suspensions were prepared using mechanical tissue dissociation, followed by ACK-mediated erythrocyte lysis. CD19$^{+}$ B cells were isolated using CD19 MicroBeads (Miltenyi Biotec). Purity analysis via FACS revealed >90% CD19$^{+}$ B cells and <0.15% CD138$^{+}$TACI$^{+}$ plasma cells in the isolated samples. IL-6Rα$^{+/+}$ and IL-6Rα$^{-/-}$ cells were mixed at a 1:1 ratio and resuspended in PBS$^{-/-}$ without supplements. A total of 1.0 x10$^{7}$ cells in 200 µl was intravenously injected into the lateral tail vein of IL-6$^{Fp3\text{-}OE}$ and control recipient mice on mixed CD45.1/CD45.2 background. Eight, and ten days post-transfer, recipient mice were analyzed for plasma cell differentiation among IL-6Rα$^{+/+}$ and IL-6Rα$^{-/-}$ transferred cells using flow cytometry.

### Supplemental Tables

| Antigen | Fluorochrome | Clone | Supplier |
|---|---|---|---|
| CD11b | BV510 | M1/70 | BioLegend |
| CD11b | BV711 | M1/70 | BioLegend |
| CD11c | RB780 | HL3 | BD Biosciences |
| CD126 (IL-6Rα) | BV421 | D7715A7 | BD Biosciences |
| CD138 | BB700 | 281-2 | BD Biosciences |
| CD138 | BV421 | 281-2 | BioLegend |
| CD138 | PE | 281-2 | BD Biosciences |
| CD19 | Biotin | 6D5 | BioLegend |
| CD19 | PE-Cy7 | 6D5 | BioLegend |
| CD19 | APC-Cy7 | 6D5 | BioLegend |
| CD19 | BUV563 | 1D3 | BD Biosciences |
| CD19 | BUV395 | 1D3 | BD Biosciences |
| CD267 (TACI) | PE | 8F10 | BioLegend |
| CD267 (TACI) | BV421 | 8F10 | BD Biosciences |
| CD4 | BUV395 | RM4-5 | BD Biosciences |
| CD4 | BV510 | RM4-5 | BioLegend |
| CD45 | BV510 | 30-F11 | BioLegend |
| CD45 | BUV805 | 30-F11 | BD Biosciences |
| CD45.1 | FITC | A20 | BioLegend |
| CD45.1 | PE-Cy7 | A20 | BioLegend |
| CD45.2 | APC | 104 | eBioscience |
| CD45.2 | FITC | 104 | eBioscience |
| CD45R (B220) | PerCP | RA3-6B2 | BioLegend |
| CD45R (B220) | BV786 | RA3-6B2 | BD Biosciences |
| CD80 | RB705 | 16-10A1 | BD Biosciences |
| Fixable Viability Dye eFluor™ 506 | | | eBioscience |
| Fixable Viability Dye eFluor™ 780 | | | eBioscience |
| Fixable Viability Stain 440UV | | | BD Biosciences |
| Foxp3 | APC | FJK-16s | eBioscience |
| GFP | PerCP-Cy5.5 | FM264G | BioLegend |
| Ig κ light chain | PE-Cy7 | 187.1 | BD Biosciences |
| Ig λ light chain | Biotin | polyclonal | Southern Biotech |
| IgD | FITC | 11-26c | eBioscience |
| IgG2a, κ (isotype control) | PE | MOPC-173 | BD Biosciences |
| IgG2b, κ (isotype control) | BV421 | R35-38 | BD Biosciences |
| IgM | FITC | II/41 | eBioscience |
| IgM | Biotin | II/41 | eBioscience |
| Ki67 | RY703 | B56 | BD Biosciences |
| Ly6C | V450 | HK1.4 | BD Biosciences |

| | | | |
|---|---|---|---|
| Ly6C | PerCP | HK1.4 | BioLegend |
| Ly6C | BV570 | HK1.4 | BioLegend |
| Ly6G | FITC | 1A8 | BioLegend |
| Ly6G | PE | 1A8 | BioLegend |
| MHCII | APC-H7 | M5/114.15.2 | BD Biosciences |
| NK1.1 | SparkRed718 | S17016D | BioLegend |
| RORγt | RB613 | Q31-378 | BD Biosciences |
| STAT3 (py705) | PE | 4/P-STAT3 | BD Biosciences |
| Streptavidin | PerCP | | BioLegend |
| Streptavidin | FITC | | eBioscience |
| Streptavidin | BV421 | | BioLegend |
| Streptavidin | BUV563 | | BD Biosciences |
| TCRβ | FITC | H57-597 | BioLegend |
| TCRβ | PE-Cy7 | H57-597 | BioLegend |
| TCRβ | SuperBright780 | H57-597 | eBioscience |
| TCRβ | BV786 | H57-597 | BD Biosciences |
| PAX-5 | PE | 1H9 | Biolegend |
| IRF4 | AFl647 | 3E4 | H57-597 |

**Supplemental Table 1. Antibodies, streptavidin conjugates, and fluorescent dyes used for flow cytometry analysis.**

### Supplemental Figures

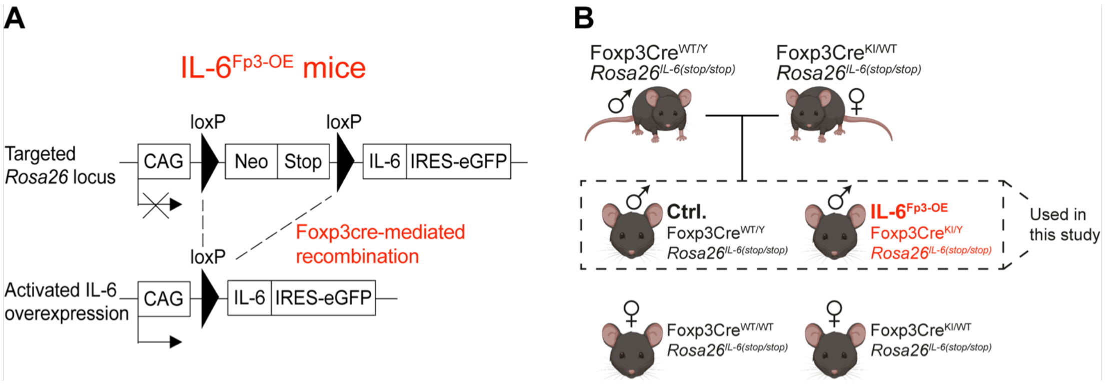


**Supplemental Figure 1. Generation of IL-6$^{Fp3\text{-}OE}$ mice.**
**A** – Genetic construct inserted into the Rosa26 locus to enable IL-6 overexpression in a Foxp3-Cre-dependent manner. The Cre recombinase under the control of the Foxp3 promoter (Foxp3Cre) used in the current study is inserted into the endogenous Foxp3 locus on the X chromosome.
**B** – Breeding strategy used in the current study. To generate experimental (IL-6$^{Fp3\text{-}OE}$) and control (Ctrl.) littermate mice, sires homozygous for the *Il6* transgene were mated with dams homozygous for the *Il6* transgene and heterozygous for the Foxp3-Cre transgene. Offspring males homozygous for the *Il6* transgene and hemizygous for Foxp3-Cre (IL-6$^{Fp3\text{-}OE}$) were used as experimental animals. Their male littermates without Foxp3-Cre were used as controls (Ctrl.) This breeding scheme allowed maintenance of breeder mice without phenotypic burden and maximized the generation of experimental and control littermates.
CAG – chicken β-actin promoter; Neo – neomycin resistance cassette; Stop – transcriptional stop cassette; IRES – internal ribosome entry site; eGFP – enhanced green fluorescent protein; WT – wild type allele; Y – Y chromosome; KI – knock-in allele; Fp3 – Foxp3-Cre; OE – overexpression.

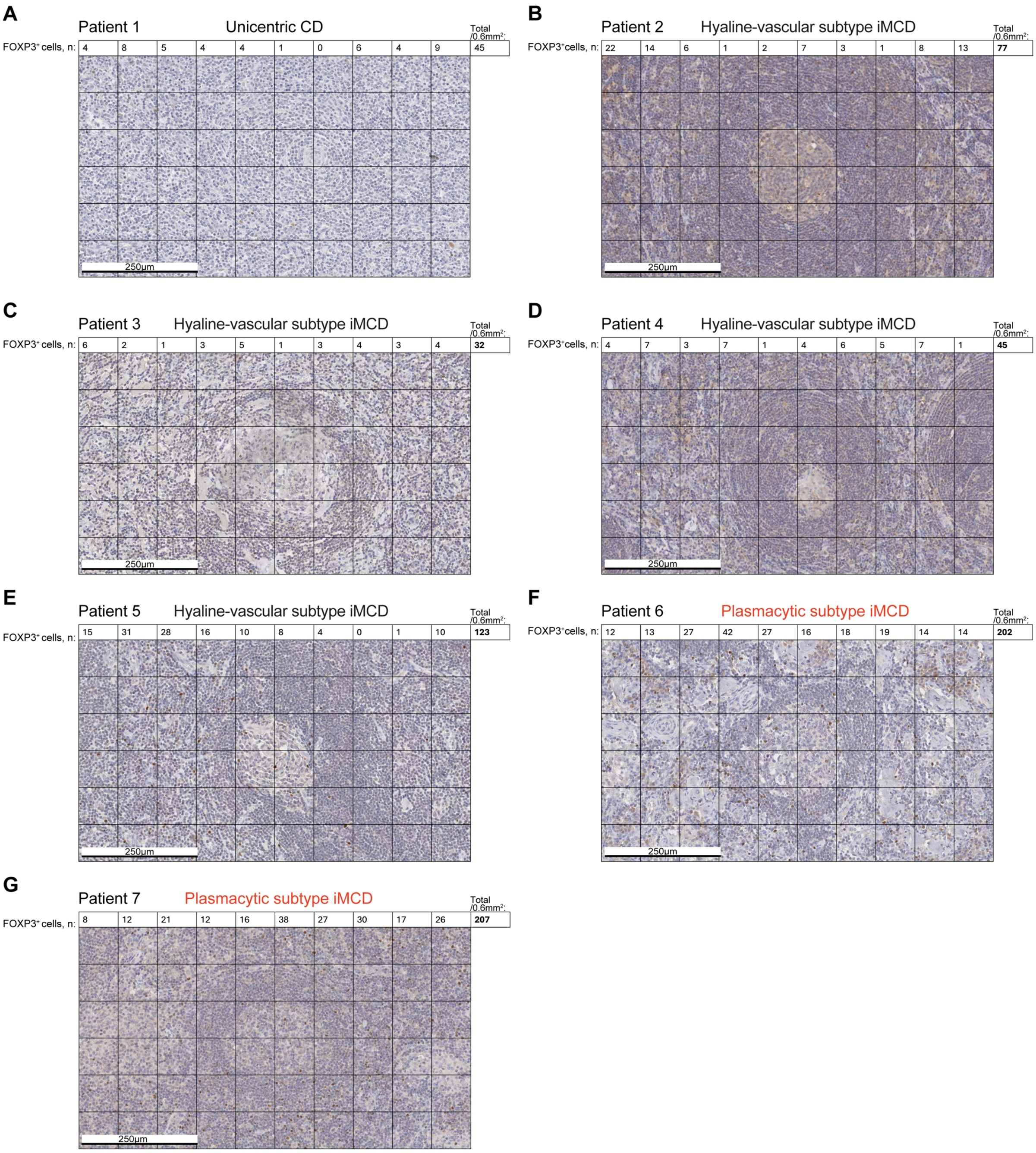


**Supplemental Figure 2. Analysis of FOXP3 expression in lymph nodes of patients with distinct Castleman disease subtypes.**

**A** – Histological analysis of a jaw angle and parotid lymph node (LN).
**B** – Histological analysis of a jaw angle LN.
**C** – Histological analysis of a supraclavicular LN.
**D** – Histological analysis of an LN from an unknown location.
**E** – Histological analysis of a retroperitoneal LN.
**F** – Histological analysis of a groin LN.
**G** – Histological analysis of a neck block LN.

Castleman disease subtypes analyzed: unicentric CD (A), hyaline-vascular iMCD (B, C, D, E), and plasmacytic iMCD (F and G). Each LN corresponds to an individual patient (1-7). Histological sections were stained with hematoxylin and anti-FOXP3 antibodies (brown staining). Images were acquired at 40× magnification using a light microscope. Scale bars indicate 250 µm. Sections were divided into quadrants, and FOXP3-positive cells were manually counted and quantified per 0.6 mm².

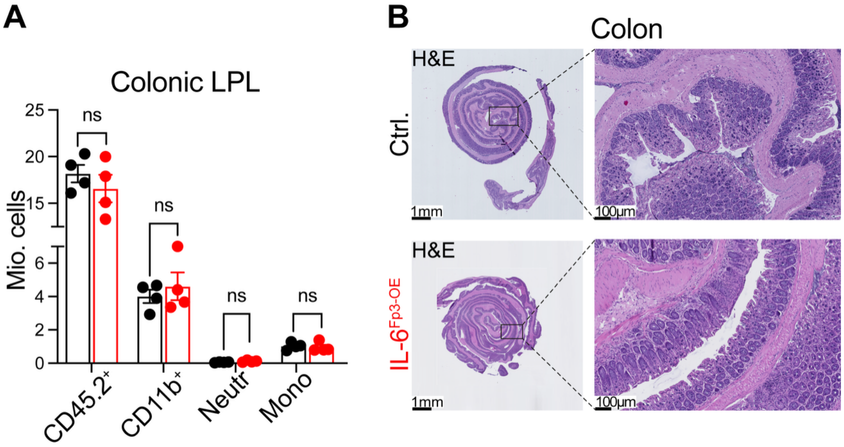


**Supplemental Figure 3. Absence of gut-directed autoimmunity in IL-6$^{Fp3\text{-}OE}$ mice.**
**A** – Absolute numbers of immune cells in the colonic lamina propria (LPL) based on FACS analysis. n=4 mice per genotype, 8 weeks old. Neutr. – Neutrophilic granulocytes (gated on CD45.2$^{+}$ TCRβ$^{-}$ CD19$^{-}$ CD11b$^{+}$ Ly6G$^{hi}$ Ly6C$^{int}$). Mono – Monocytes (gated on CD45$^{+}$ TCRβ$^{-}$ CD19$^{-}$ CD11b$^{+}$ Ly6G$^{neg}$ Ly6C$^{int\text{-}hi}$).
**B** – Histological analysis (H&E staining) of the colon. n=3 mice per genotype, 8 weeks old.
Graph (A) shows mean ± SEM. Each dot represents an individual mouse. P values were calculated using unpaired Student's t-test (A) and are indicated as follows: ns – not significant ($p > 0.05$).

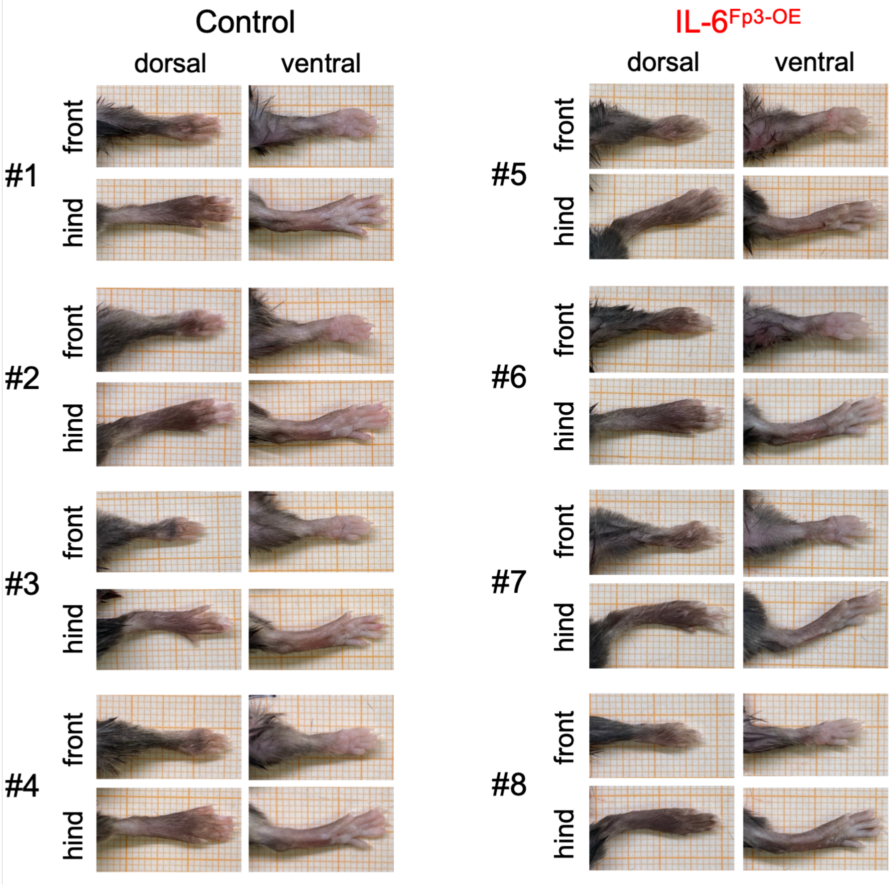


**Supplemental Figure 4. Absence of joint swelling in IL-6$^{Fp3\text{-}OE}$ mice.**
Macroscopic examination of the front and hind paws of 9-10week-old mice, shown from dorsal and ventral views. Images were acquired against a 1-mm grid background. Representative pictures of n=4 of Control, n=4 of IL-6$^{Fp3\text{-}OE}$ mice.

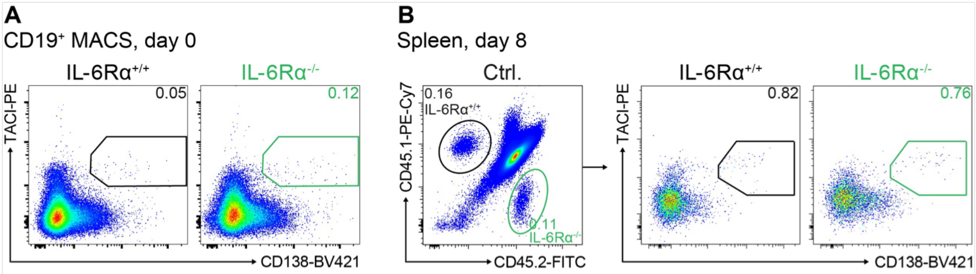


**Supplemental Figure 5. Adoptive co-transfer of IL-6Rα$^{+/+}$ and IL-6Rα$^{-/-}$ B cells.**

**A** – FACS analysis showing the near absence of CD138$^{+}$TACI$^{+}$ plasma cells in the CD19$^{+}$ MACS-isolated cell preparations prior to transfer into recipient mice.

**B** – FACS analysis of plasma cell differentiation in the spleen of naïve wild-type recipient mice, 8 days post-transfer.

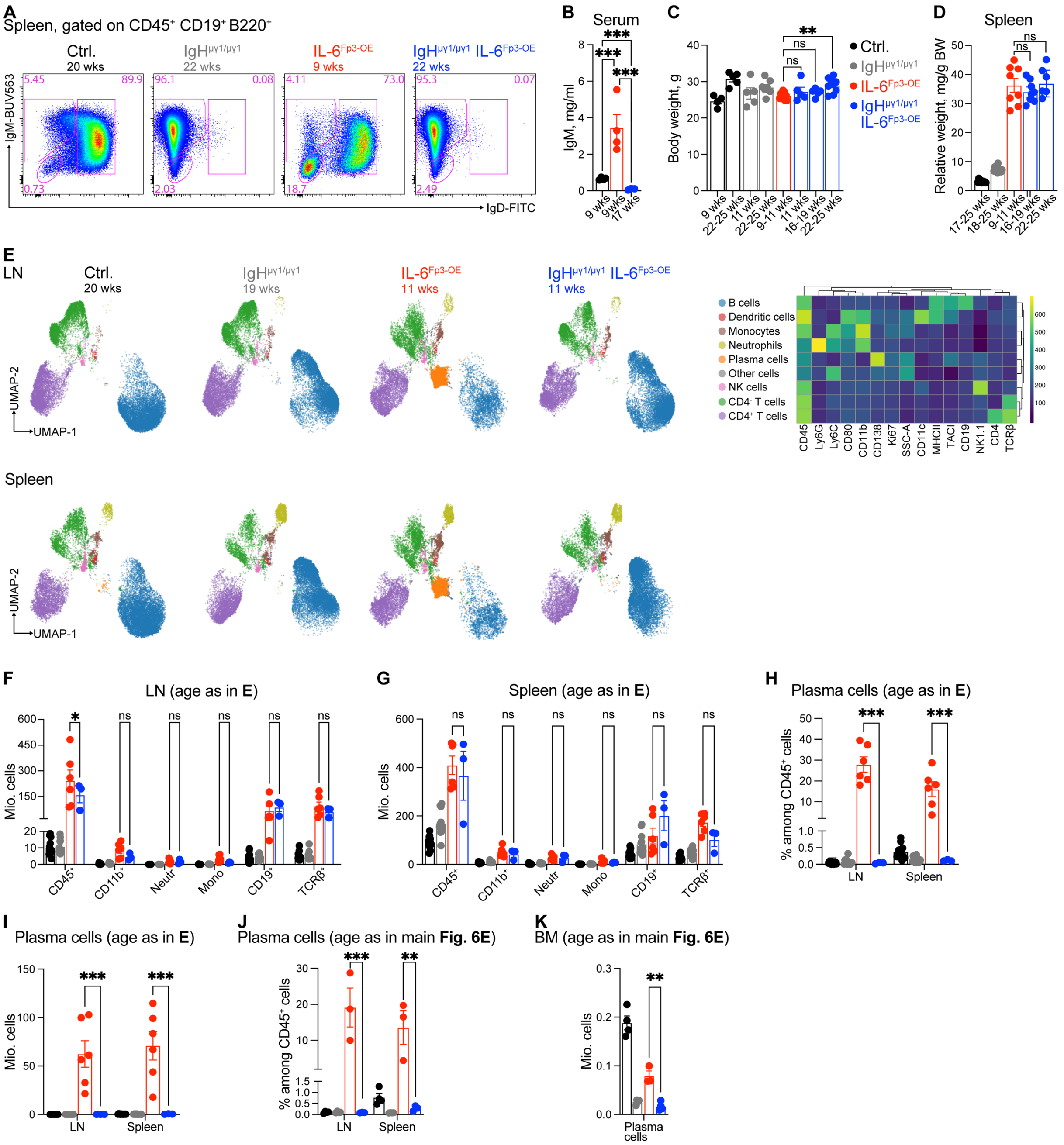


**Supplemental Figure 6. IgH$^{\mu\gamma1/\mu\gamma1}$ background protects IL-6$^{Fp3\text{-}OE}$ mice from plasmacytic iMCD-like disease.**

**A** – FACS analysis of surface IgM and IgD expression on CD19$^{+}$B220$^{+}$ B cells in the spleen of mice with the indicated age and genotype.

**B** – IgM levels in the serum of mice with the indicated age and genotype measured by ELISA.

**C** – Body weight measurements of mice with the indicated age and genotype.

**D** –Spleen weight analysis of mice with the indicated age and genotype. Relative spleen weight was calculated by dividing spleen weight (mg) by body weight (g). BW – body weight.

**E** – High-dimensional flow cytometric analysis of immune cell populations (gated on live CD45$^{+}$ cells) in lymph nodes (LN) of mice with the indicated age and genotype. The heat map displays respective individual clusters with median expression profiles identified by FlowSOM algorithm. A total of 20,000 cells per sample were used to generate UMAP. An overlay of the clusters identified by FlowSOM is shown.

**F** – FACS analysis of immune cell composition in lymph nodes of mice. Neutr. – Neutrophilic granulocytes (defined as CD45$^{+}$ TCRβ$^{-}$ CD19$^{-}$ CD11b$^{+}$ Ly6G$^{hi}$ Ly6C$^{int}$). Mono – Monocytes (defined as CD45$^{+}$ TCRβ$^{-}$ CD19$^{-}$ CD11b$^{+}$ Ly6G$^{neg}$ Ly6C$^{int\text{-}hi}$).

**G** – FACS analysis of immune cell composition in the spleen of mice. Neutr. and mono., defined as in **E**.
**H** – Frequency of $CD138^{+}TACI^{+}$ plasma cells in the lymph nodes and spleen
**I** – Number of $CD138^{+}TACI^{+}$ plasma cells in the lymph nodes and spleen
**J** – Frequency of $CD138^{+}TACI^{+}$ plasma cells in the lymph nodes and spleen of 20-week-old control mice, 22-24-week-old $IgH^{\mu\gamma1/\mu\gamma1}$ mice, 8-9-week-old IL-$6^{Fp3\text{-}OE}$, and 22-week-old $IgH^{\mu\gamma1/\mu\gamma1}$ IL-$6^{Fp3\text{-}OE}$ mice.
**K** – Number of $CD138^{+}TACI^{+}$ plasma cells in the bone marrow (two femora and one tibia) of 20-week-old control mice, 22-24-week-old $IgH^{\mu\gamma1/\mu\gamma1}$ mice, 8-9-week-old IL-$6^{Fp3\text{-}OE}$, and 22-week-old $IgH^{\mu\gamma1/\mu\gamma1}$ IL-$6^{Fp3\text{-}OE}$ mice.
Graphs (C, D, F, G, H, I, J, K) show mean ± SEM. Each dot represents an individual mouse. P values were calculated using unpaired Student's t-test (C, D, F, G, H, I, J, K) and are indicated as follows: * ($p < 0.05$), ** ($p < 0.01$), *** ($p < 0.001$); ns – not significant ($p > 0.05$).

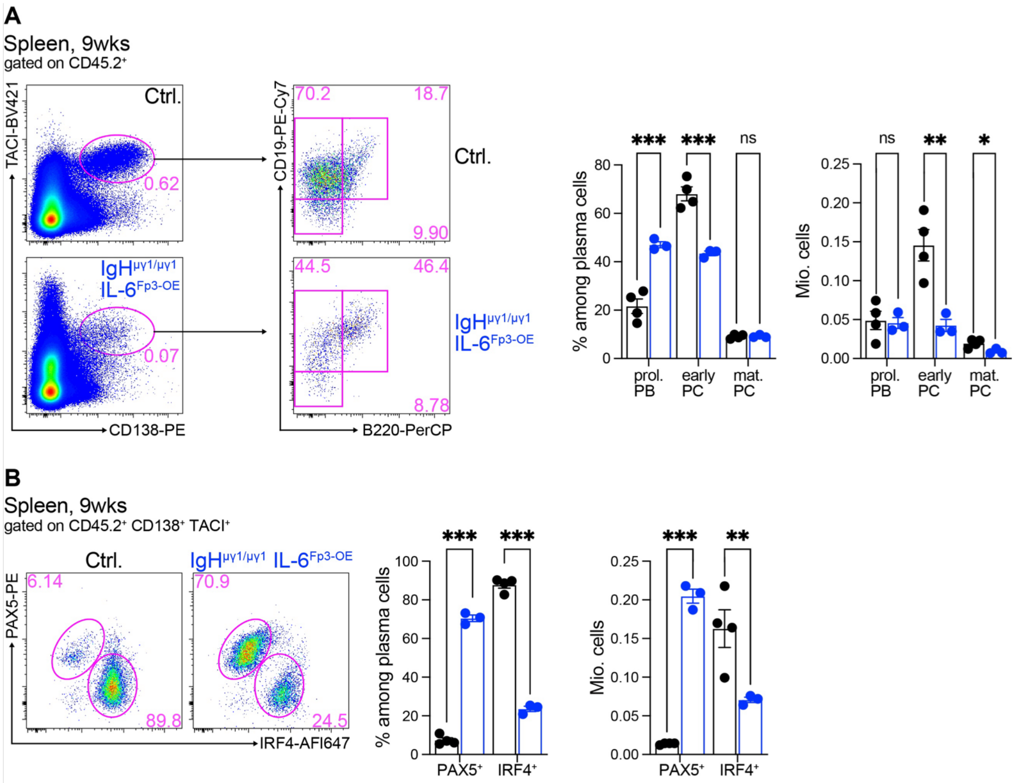


**Supplemental Figure 7. Analysis of plasma cells development in IgH$^{\mu\gamma1/\mu\gamma1}$ IL-6$^{Fp3\text{-}OE}$ mice.**
**A** – FACS analysis of CD138$^{+}$TACI$^{+}$ cells and their subsets of 9-week-old control mice and 9-week-old IgH$^{\mu\gamma1/\mu\gamma1}$ IL-6$^{Fp3\text{-}OE}$ mice. Proliferating plasmablasts (prol. PB) were defined as CD19$^{int}$B220$^{int}$, early plasma cells (early PC) as CD19$^{int}$B220$^{lo}$, and mature plasma cells (mat. PC) as CD19$^{neg\text{-}lo}$B220$^{neg\text{-}lo}$.
**B** – FACS analysis of intracellular PAX5 and IRF4 expression of CD138$^{+}$TACI$^{+}$ cells isolated from 9-week-old control mice and 9-week-old IgH$^{\mu\gamma1/\mu\gamma1}$ IL-6$^{Fp3\text{-}OE}$ mice in the spleen.
Graphs (A, B) show mean ± SEM. Each dot represents an individual mouse. P values were calculated using unpaired Student's t-test (A, B) and are indicated as follows: * ($p < 0.05$), ** ($p < 0.01$), *** ($p < 0.001$); ns – not significant ($p > 0.05$).

### Supplemental References


1. Mufazalov IA, Andruszewski D, Schelmbauer C, et al. Cutting Edge: IL-6-Driven Immune Dysregulation Is Strictly Dependent on IL-6R alpha-Chain Expression. *J Immunol*. 2020;204(4):747-751.
2. Knopp T, Jung R, Wild J, et al. Myeloid cell-derived interleukin-6 induces vascular dysfunction and vascular and systemic inflammation. *Eur Heart J Open*. 2024;4(4):oeae046.
3. Wing K, Onishi Y, Prieto-Martin P, et al. CTLA-4 control over Foxp3+ regulatory T cell function. *Science*. 2008;322(5899):271-275.
4. Waisman A, Kraus M, Seagal J, et al. IgG1 B cell receptor signaling is inhibited by CD22 and promotes the development of B cells whose survival is less dependent on Ig alpha/beta. *J Exp Med*. 2007;204(4):747-758.
5. Wunderlich FT, Strohle P, Konner AC, et al. Interleukin-6 signaling in liver-parenchymal cells suppresses hepatic inflammation and improves systemic insulin action. *Cell Metab*. 2010;12(3):237-249.
6. Caton ML, Smith-Raska MR, Reizis B. Notch-RBP-J signaling controls the homeostasis of CD8- dendritic cells in the spleen. *J Exp Med*. 2007;204(7):1653-1664.
7. Van Gassen S, Callebaut B, Van Helden MJ, et al. FlowSOM: Using self-organizing maps for visualization and interpretation of cytometry data. *Cytometry A*. 2015;87(7):636-645.
8. McInnes L, Healy J, Melville J. Umap: Uniform manifold approximation and projection for dimension reduction. *arXiv preprint arXiv:180203426*. 2018.
9. Wickham H. ggplot2 : Elegant Graphics for Data Analysis. Use R!,. Cham: Springer International Publishing : Imprint: Springer,; 2016:1 online resource (XVI, 260 pages 232 illustrations, 140 illustrations in color.
10. Kolde R. Pheatmap: pretty heatmaps. *R package version*. 2019;1(2):726.
